# Highly task-specific and distributed neural connectivity in working memory revealed by single-trial decoding in mice and humans

**DOI:** 10.1101/2021.04.20.440621

**Authors:** Daniel Strahnen, Sampath K.T. Kapanaiah, Alexei M. Bygrave, Birgit Liss, David M. Bannerman, Thomas Akam, Benjamin F. Grewe, Elizabeth L. Johnson, Dennis Kätzel

## Abstract

Working memory (WM), the capacity to briefly and intentionally maintain mental items, is key to successful goal-directed behaviour and impaired in a range of psychiatric disorders. To date, several brain regions, connections, and types of neural activity have been correlatively associated with WM performance. However, no unifying framework to integrate these findings exits, as the degree of their species- and task-specificity remains unclear. Here, we investigate WM correlates in three task paradigms each in mice and humans, with simultaneous multi-site electrophysiological recordings. We developed a machine learning-based approach to decode WM-mediated choices in individual trials across subjects from hundreds of electrophysiological measures of neural connectivity with up to 90% prediction accuracy. Relying on predictive power as indicator of correlates of psychological functions, we unveiled a large number of task phase-specific WM-related connectivity from analysis of predictor weights in an unbiased manner. Only a few common connectivity patterns emerged across tasks. In rodents, these were thalamus-prefrontal cortex delta- and beta-frequency connectivity during memory encoding and maintenance, respectively, and hippocampal-prefrontal delta- and theta-range coupling during retrieval, in rodents. In humans, task-independent WM correlates were exclusively in the gamma-band. Mostly, however, the predictive activity patterns were unexpectedly specific to each task and always widely distributed across brain regions. Our results suggest that individual tasks cannot be used to uncover generic physiological correlates of the psychological construct termed WM and call for a new conceptualization of this cognitive domain in translational psychiatry.

## Introduction

Working memory (WM) refers to the capacity to store and manipulate contents of perception and thought at the forefront of attention over seconds to minutes^1, 2^. Several psychiatric and neurological disorders are characterised by severe and pharmaco-resistant impairments of WM^3^. Studies in humans, non-human primates, and rodents^4^ have centrally implicated the prefrontal cortex (PFC)^5–12^, parietal cortex^11, 13^, medio-dorsal thalamus (MD)^14, 15^, and hippocampus^8, 16–21^ in WM processes. In particular, connectivity between these brain structures correlates with the performance in specific tasks^11, 22–24^ (see Supplementary Table 1 for an overview of rodent studies).

However, it remains elusive which kinds of neural connectivity actually mediate WM. Several-partially conflicting – reports suggest that specific brain regions, frequency bands^25–27^ and types of inter-regional coupling (or metrics to analyse) are essential for WM. Studies in rodents, for example, have implicated network oscillations in either the delta (δ)^28^, theta (θ)^27, 29, 30^, beta (β)^14, 26^, or gamma (γ)^25^ frequency, in coupling between such frequency bands (esp. θ−γ)^31^ or between oscillations and local spiking of neurons^14, 27, 31–33^, or in interactions between either the ventral (vHC)^31, 33^ or dorsal hippocampus (dHC)^26, 29, 34^ and PFC. On a T-maze rewarded-alternation task, both vHC➔PFC connectivity^33^ and dHC-PFC coupling via the thalamic nucleus reuniens^34^ have been claimed to mediate the encoding of WM-contents using optogenetic inhibition experiments. In addition to this diversity of findings in rodents - which mainly originate from recordings in spatial alternation tasks (Supplementary Table 1) - it remains largely unknown to what extent the discovered WM substrates generalize across different tasks and species.

Moreover, it remains unclear if this diversity of reported associations between specific neural activities and WM performance implicates that, indeed, several types of connectivity and connections are engaged in WM simultaneously. Alternatively, it could be due to differences between task paradigms, species, specific rodent models, or analysis procedures. Several factors may explain the current uncertainty regarding the physiological correlates of WM. This includes the difficulty of separating correlates of more basic functions like motivation, spatial processing, and attention from the actual WM component of the task - especially in rewarded-alternation assays that are mostly used in rodents (Supplementary Table 1)^35^. Further uncertainty arises from the typical analytical approach of correlating within-subject averages of performance (WM accuracy score)^26, 29, 31^, trial type (correct vs. incorrect)^33^, or trial phase (sample vs. choice phase)^30, 33^ with within-subject averages of a select connectivity measure. Such correlations do not necessitate mechanistic causation but could be indirect or even epiphenomenal. Finally, the *streetlight effect* has recently been highlighted as a rather principal limitation of studies of physiological correlates of psychological function^36^. In this context, this term denotes the missing of neurophysiological correlates due to a biased selection of only few investigated brain regions, connections, or activity parameters. We recently showed that the rich plethora of widely used functional connectivity measures indeed contains a considerable number of non-redundant metrics^37^, which entails the possibility for seemingly contradictory findings and inconclusive null results.

To identify inter-regional neural connectivity that is associated with WM in mice and humans, we addressed these issues in several ways: First, we comparatively assessed three distinct visual WM tasks in each species. In mice, we applied an operant 5-choice delayed-*matching*-to-sample (DMTS) task as well as a T-maze-based and an operant 2-choice delayed-*non*-matching-to-sample (DNMTS) task. In humans, we analysed an existing dataset where three distinct task types which featured the same temporal structure and were inter-mixed within one session^20^. Therefore, in each species, the non-WM related differences between the applied tasks were relatively small, which benefits an optimal comparison of the neurophysiological basis of different types of WM. Second, during these tasks, local field potentials (LFPs) were simultaneously recorded from four sites (PFC, dHC, vHC, and MD thalamus) in rodents, and from three sites (PFC, medial temporal lobe, MTL, and orbitofrontal cortex, OFC) in humans. Third, to maximize connectivity-related information obtained from these multi-site recordings, we extracted a large set of measures of inter-regional neural coupling. The composition of this set was optimized based on our prior analysis of redundancies between the most commonly used LFP-based coupling metrics^37^. Each metric was determined in three or four task phases, four or five frequency bands, and as both absolute and relative measures. This yielded a total of 960 and 1344 connectivity measures in mice and humans, respectively, in addition to ∼240 measures of local activity. Finally and most importantly, we applied machine learning (ML) to predict WM choices on a single-trial basis from these high-dimensional patterns of functional connectivity. Data-driven approaches have recently been deployed to reduce the *streetlight effect*^36, 38–40^. These studies have applied *unsupervised* classification procedures to extract electrophysiological correlates of stress responses^39^, depression vulnerability^38^, and anxiety^36^ from connectivity patterns obtained from multi-site LFP-recordings in mice, collectively termed the *electome*. To establish a largely unbiased search for WM correlates within high-dimensional electome patterns, in contrast, we translated the supervised, trial-based nature of WM tasks (pre-defining choices as correct or incorrect) into our analysis^10, 21, 24, 33, 41, 42^. Implementing a *supervised* decoding analysis allowed us to harness the power of prediction to prove the presence of behaviourally relevant information in complex patterns of neural signals^43^. The possibility of trial-by-trial prediction of WM-based choices from the spiking activity of large ensembles of neurons in rodent PFC^10, 33^, monkey inferior temporal cortex^41^, or human hippocampus^21^ has recently been demonstrated. In these attempts, however, prediction was done with classifiers that were trained specifically for a single animal and task session, since the predictor variables were individual neurons. In contrast, we applied ML across datasets from multiple subjects and sessions and used connectivity and activity metrics from multiple sites as predictor variables. In this way, subsequent quantification of the predictive power of each electrophysiological variable allowed a largely unbiased identification of connectivity patterns associated with WM in mice and humans as verified by trial-by-trial predictive power.

## Results

### Correct choices in DMTS working memory are associated with distinct signatures of connectivity

To evaluate associations between WM and electrophysiological measures of inter-regional connectivity, we first implanted mice with chronic field electrodes in PFC, MD, vHC, and dHC and tested them in three spatial working memory (SWM) tasks (Fig. 1a-f): first, an operant DMTS 5-choice SWM (5-CSWM) task, subsequently the T-maze rewarded alternation task, and finally an operant DNMTS 2-choice SWM (2-CSWM) task. To allow for simultaneous LFP recordings during operant tasks, we developed a custom-designed operant box optimized for implanted and tethered animals (Fig. 1c) that is tightly integrated with electrophysiological recordings via pyControl software and microcontroller modules^44^. The set of three tasks (Fig. 1d-f) was chosen to retain comparability due to their shared visuo-spatial nature and distinct individual differences (i.e., operant DMTS vs. DNMTS; maze-based DNMTS vs. operant DNMTS). In addition, the T-maze task was included due to its wide usage (see e.g., Supplementary Table 1). The 5-CSWM was specifically designed to limit the usage non-WM mediation strategies by the animal - due the large number of choice configurations, requirement to shuttle between opposite walls of the box, and delay-periods in total darkness, as we described in detail recently^45^. Also, both operant tasks provide tight control over the timing of behavioural events and deliver intrinsic control variables for WM-enabling psychological functions like attention, measured by sample phase (SP) accuracy, and motivational drive, measured by reward latency. In each task, we applied extensions of the delay between the SP and the choice phase (CP) across which the memorized information needed to be maintained in order to strongly engage WM capacity. As expected for WM assays^2^, such delay challenges significantly decreased WM choice accuracy of the mice (Fig. 1g-i).

**Fig. 1.**
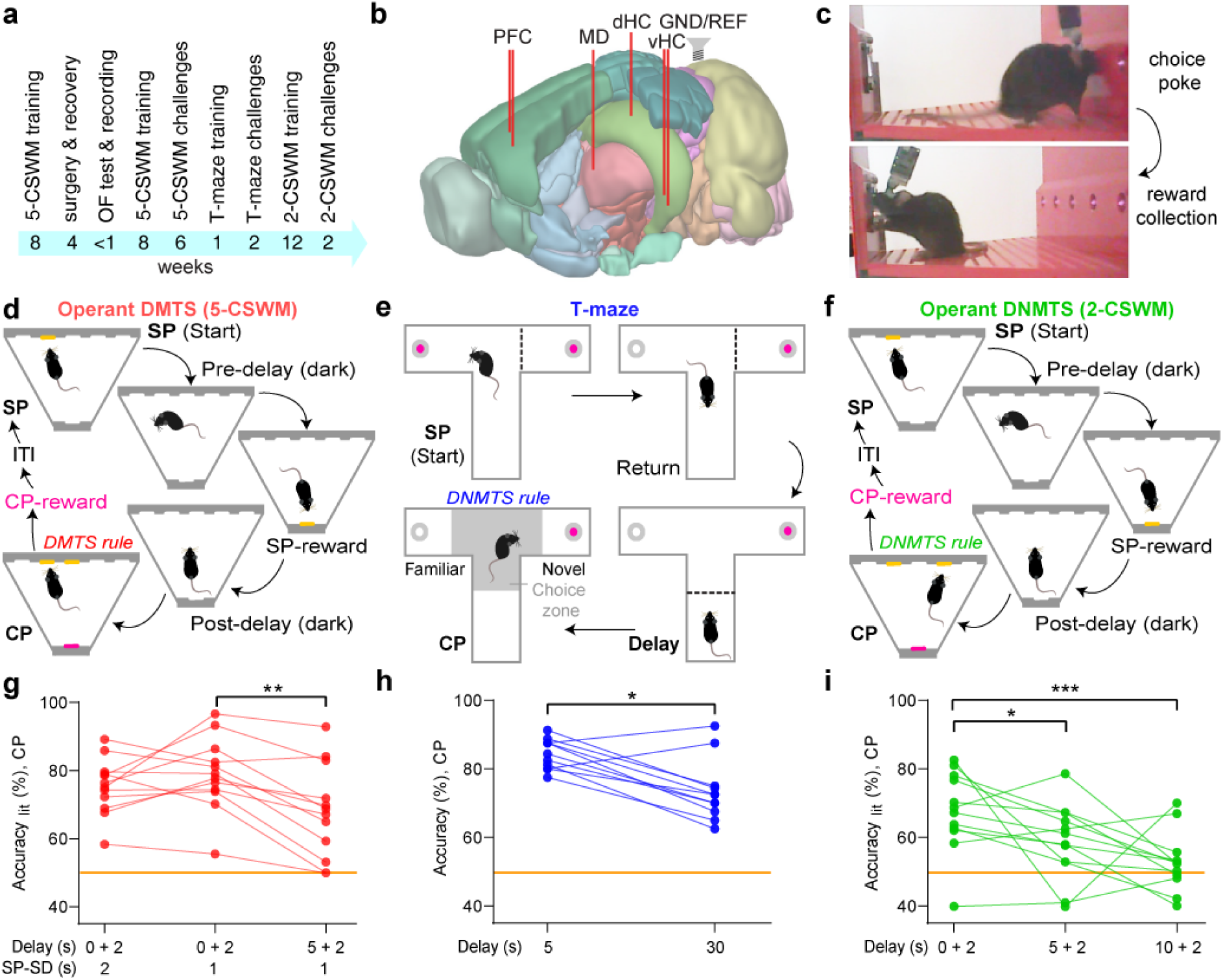
Rodent SWM tasks. (**a**) Timeline of experiments in the analysed cohort (see Methods). (**b**) Illustration of electrode placements in the mouse brain (image taken from Allen Brain Atlas); pairs of electrodes were inserted into PFC and vHC. (**c**) Lack of obstruction of tethered mice with mounted headstage during poking of choice hole (top) and reward collection (bottom) in custom-made pyControl operant boxes. (**d-f**) Illustration of DMTS 5-CSWM^45^ (d), T-maze rewarded alternation (e), and DNMTS 2-CSWM (f) tasks; in (d,f), choices in SP and CP need to be made at the 5-choice wall (top), while rewards for correct responses in each phase are collected on the opposite wall (magenta, bottom). (**e-i**) WM performance measured as response accuracy in the CP (% correct choices relative to available indicated options) in 12 mice in each set of challenge conditions including their respective baseline with simultaneous electrophysiological recordings. Delay length determining the WM challenge stated on x-axes; for operant tasks (e,j) pre-+ post-delay (referring to set delays before and after SP-reward collection) are indicated. SP-SD, stimulus duration in sample phase. Asterisks indicate differences between challenge, Sidak-post-hoc tests conducted after significant main effect of challenge, RM-ANOVA. Orange line, chance level performance. * *P* < 0.05; ** *P* < 0.01; *** *P* ≤ 0.001.

To initially explore possible WM correlates among measures of neural connectivity, we computed time-resolved spectrograms of four largely non-redundant^37^ connectivity metrics aligned to the time of correct and incorrect SP and CP responses for the distinct phases of the 5-CSWM task (*non-directed*: coherence, Coh; weighted Phase-Lag-index, wPLI; *directed*: Granger causality, GC; partial directed coherence, PDC; Supplementary Fig. 1-2). We subtracted spectra of correct SP responses or incorrect CP responses from those of correct CP responses to eliminate neural representations of poking action, execution of a reward-related response, and attention (Fig. 2a-d, Supplementary Fig. 3). This qualitative analysis suggested a complex pattern of connectivity associated with correct WM decisions, including elevated hippocampal-prefrontal connectivity in the low γ-range (30-48 Hz) immediately after the CP response (Fig. 2a-b) and sustained connectivity in the PFC➔MD and vHC➔dHC θ-range (5-12 Hz) during the delay (Fig. 2c-d).

**Fig. 2.**
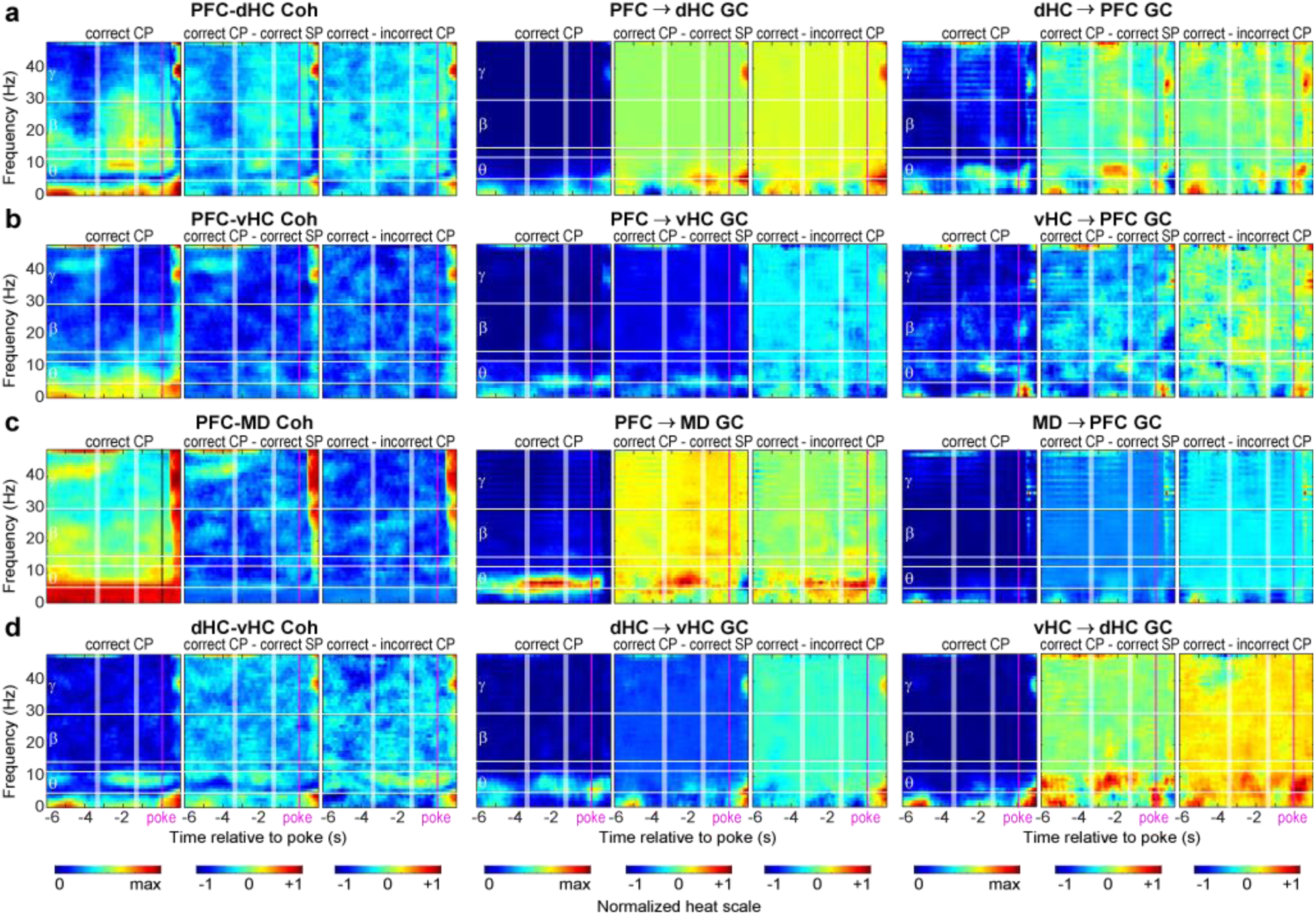
Connectivity in the DMTS 5-CSWM task. (**a-d**) Spectrograms depicting min-max normalized coherence (Coh) and Granger causality (GC) for the connections stated above each triplet panel for the delay and CP of the 5-CSWM task, temporally aligned to the choice poke entry (p, vertical magenta lines) showing 6 s before until 1 s after the choice poke (x-axes in (d)); the start and end of the post-delay shown by white stripes corresponding to mean±SD as determined by CP response latency. Each triplet shows the absolute value (left), the difference between the former and either the prior correct SP (middle), or incorrect CPs (right). Horizontal white lines show borders between analysed frequency bands, stated on the left. See Supplementary Fig. 1-2 for spectrograms of absolute values in correct and incorrect trials of the CP and SP, respectively, and Supplementary Fig. 3 for the same display as (a-d) for wPLI and PDC.

### Trial-by-trial prediction of DMTS WM-mediated choices from local and long-range neural activity

The ability to predict behavioural choices from neural activity may be regarded as evidence that such activity encodes aspects of these choices^33, 43^. We assessed individual connectivity variables as five separate connectivity metrics (Coh, wPLI, GC, PDC, and cross-regional θ−γ phase-amplitude coupling, PAC) in four task phases (SP, pre-reward delay, post-reward delay, CP; see Fig. 1d), and four frequency bands (δ, 1-4 Hz, θ, 5-12 Hz, β, 15-30 Hz, low-γ, 30-48 Hz) along 4 connections (as shown in Fig. 2a-d). In the same way, indicators of local activity (power and local θ−γ PAC) were calculated within the four involved regions (dHC, vHC, PFC, MD). Additionally, all metrics were calculated in *relative* terms by dividing their obtained value by the value of that specific metric during the inter-trial-interval (ITI) before the start of the respective trial. This resulted in 240 variables for each inter-regional connection and 56 intra-regional activity variables for each area characterizing every 5-CSWM trial. We trained subspace discriminant classifiers – which proved superior among 25 different types of linear and non-linear classifiers (Supplementary Fig. 4) - to predict WM-choice trial-by-trial using the parameters contributed by each connection or region separately for trials of the final 5-CSWM challenge (1 s SP stimulus duration, SP-SD, 5 s delay; Fig. 3a). A single decoding model was generated across all subjects and its decoding accuracy was determined by predicting trials that were not part of the training dataset, i.e. by cross-validation.

**Fig. 3.**
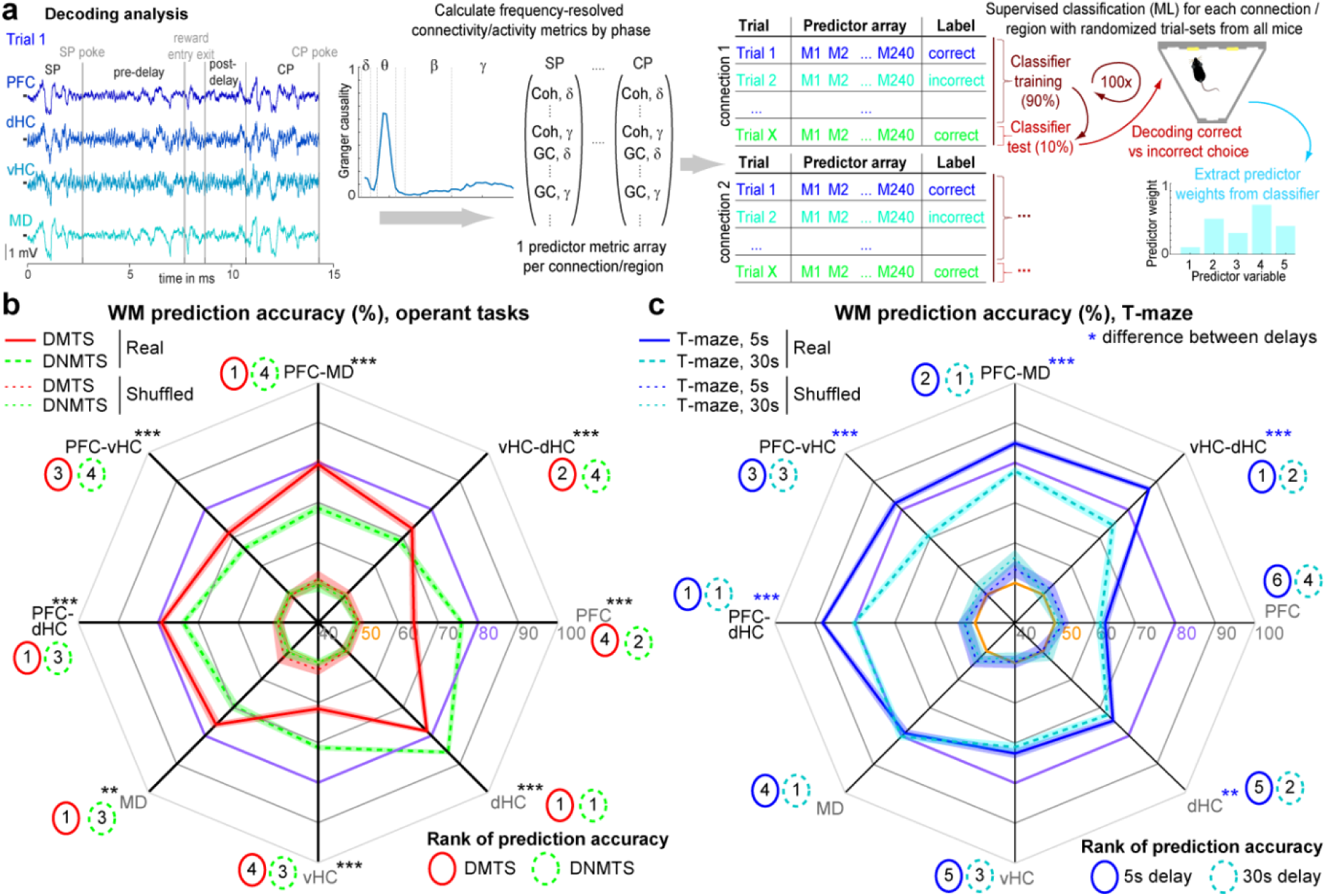
Trial-by-trial decoding of WM-based choice. (**a**) Illustration of ML-based decoding analysis (see Methods). (**b-c**) Cross-subject decoding accuracies achieved on average when using connectivity or local activity parameters of the indicated individual connections (black) or areas (grey), respectively to predict WM-based *correct* vs. *incorrect* choices in the DMTS 5-CSWM task (combined 1 s SP-SD, 5+2 s delay challenge, 2 sessions; red, b), the DNMTS 2-CSWM task (baseline, 2 sessions, green, b), the T-maze WM task with either 5s (solid blue, 4 sessions, c) or 30s (dashed blue, 4 sessions, c). Thinner dotted lines show decoding accuracies of corresponding classifiers trained with shuffled labels, remaining at chance level (50%, orange). Classifiers trained with real labels perform better than those trained with shuffled labels in all cases (*P* < 10^-17^, *t*-tests, not indicated). The accuracy of 80% is coloured in purple to aid comparison. Numbers in coloured ovals indicate the rank of the prediction accuracies achieved on average by using data from the respective connection or region. Ranks have been generated from pairwise comparisons with Tukey post-doc tests conducted after significant effects of connection/region in one-way ANOVAs (*P* < 0.0001 in all cases); connections/regions that were not significantly different from each other were assigned the same rank. Black stars in (b) indicate differences of accuracy values achieved in the two operant tasks (Tukey post-hoc tests after significant effect of task-type in ANOVA across all three tasks). Blue stars in (c) indicate pairwise differences between the two delays (uncorrected *t*-tests). ** *P* < 0.01; *** *P* ≤ 0.001. Shaded regions around mean show s.e.m. across 100 classifiers generated for each task and connection or region.

We found that individual 5-CSWM choices could be decoded with 79.4% and 79.8% maximum average accuracy when using measures of neural connectivity along the PFC-MD or the PFC-dHC connection as predictors, respectively (Fig. 3b). Using one-way ANOVA and pairwise Tukey post-hoc tests, we established a hierarchy between decoding accuracies obtained from each of the four connections and four regions. Decoding accuracies obtained from PFC-MD, PFC-dHC, local dHC and MD activity did not differ from each other and were superior to the remainder (Fig. 3b). Even though decoding accuracies varied by connection and region, they were always significantly better than those of control classifiers trained with shuffled labels (*P* < 10^-17^, *t*-tests) which, in turn, decoded indistinguishable from the 50% chance level on average (Fig. 3b). To evaluate the generality of the obtained classifiers, we assessed if they could also decode WM-based choices in data from other 5-CSWM challenge protocols. Even though the resulting decoding accuracies were generally lower compared to those achieved with data from the same protocol, they were still significantly higher than those of classifiers trained with shuffled labels (Supplementary Fig. 5). These analyses reveal that WM-based choice is encoded in LFP-based connectivity and activity measures in individual trials and that such information is widely distributed across multiple brain regions.

### Specific connections and regions are engaged differently in distinct rodent WM tasks

To investigate if this conclusion applies generally to rodent WM, we repeated the same analysis for the operant DNMTS data (final baseline sessions, 2 s delay). In this case, however, we obtained the maximum average prediction accuracy (86.1 %) from local dHC activity, rather than PFC-MD (66.1%, lowest rank of all classifiers) or PFC-dHC (73.5%) connections. Generally, in this task, local activities allowed relatively high decoding accuracies (72-77% for PFC, MD, and vHC), while coupling metrics were significantly less predictive (66-68%, *P* < 0.001, Tukey; except for dHC-PFC, Fig. 3b).

In reverse, trial-by-trial decoding of T-maze data achieved the highest average accuracies (82-88%) when using connectivity data from either one of the four connections (with dHC-connections being most predictive), whereas local activities were significantly less predictive (62-79%; *P* < 0.001, Tukey, Fig. 3c). However, decoding accuracies for information from all 4 connections decreased when analysing data from the 30 s delay challenge, in which these animals also showed lower behavioural performance (Fig. 1h, 3c), suggesting that not only task type but also task difficulty affect the information encoded in each connection.

### Specific connections and regions are engaged differently in distinct phases of the rodent DMTS 5-CSWM task

The prior analyses entail at least two conclusions: Firstly, WM-related information in a single trial is not encoded in any single region or connection, although some of them bear higher predictive power regarding WM-choice than others. Secondly, the predictive power of a given region or connection is not uniform but strongly depends on the type of WM task, indicating that different mechanisms and regions are engaged to solve distinct behavioural demands. These conclusions re-emphasize the question as to what extent oscillatory processes in distinct frequency bands, of a distinct biological type, or in a specific task phase (encoding, delay, choice) can be regarded as correlates of WM (Supplementary Table 1).

To answer this question, we took advantage of the fact that a linear classifier reveals the predictive power of each involved predictor variable according to its assigned weight. We performed Bonferroni-adjusted *t*-tests comparing the weights for each connectivity variable with the weights assigned by the classifiers trained on label-shuffled control data, and, additionally, conducted *t*-tests comparing the amplitudes of each variable between correct and incorrect trials that contributed to the classifiers. Variables for which both *t*-tests were significant were considered as bearing WM-related information (indicated by colour in Fig. 4a). This analysis revealed a relatively small set of consistent WM-related feature-classes as correlates of DMTS 5-CSWM, a majority of them in the γ-range (Fig. 4b): (*1*) PFC➔MD δ- and γ-range connectivity in the *SP*, (*2*) MD➔PFC β- and γ-range as well as dHC-vHC θ−range connectivity in the *delay*, (*3*) dHC➔PFC and MD-PFC γ-range as well as vHC-PFC δ−θ-range coupling in the *CP,* and (*4*) intra-hippocampal δ-connectivity in *all phases* (Fig. 4c-d; see also 5c).

**Fig. 4.**
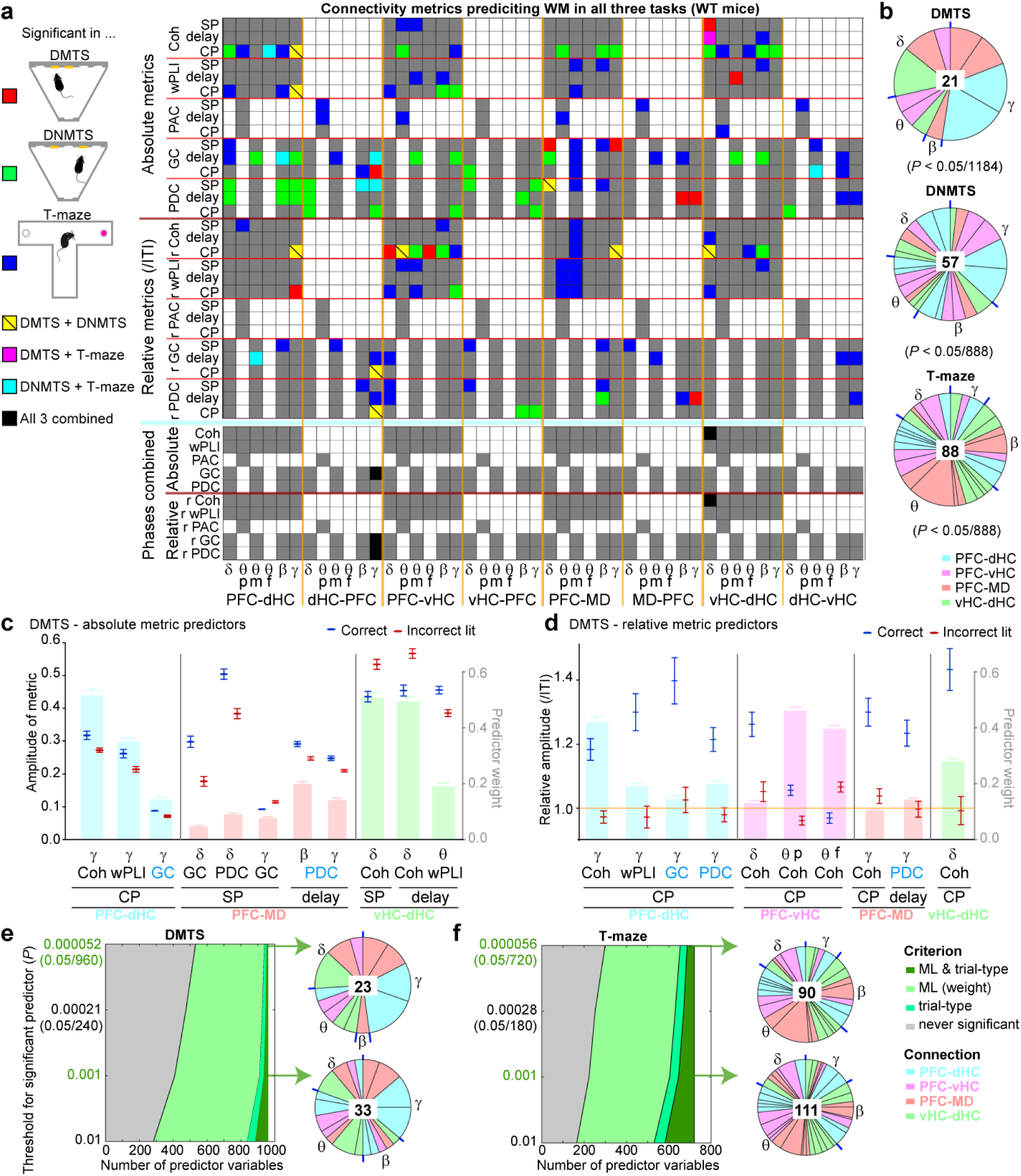
Individual connectivity measures predicting WM choice in mice. (**a**) Matrix showing all connectivity predictor variables that contributed to the connection-based classifiers shown in Fig. 3b,c. Variables that were significantly associated with WM-performance according to both their prediction weight and differences between correct and incorrect CP (*t*-tests, Bonferroni-adjusted for total number of variables, see (b)) in any of the three WM tasks are indicated by the corresponding colour, remainder in grey (white squares have no corresponding variable). Variables from the pre- and post-delay in the 5-CSWM are combined in single lines. For θ, mean amplitude (m), peak amplitude (p) and frequency of peak (f) are shown, while for all other frequency bands only the mean amplitude is used due to the absence of a clear singular peak. At the bottom, all task phases are combined and only connectivity metrics that are predictive in all three tasks (in at least one phase) are indicated in black. (**b**) Share of each connection (coded by colour) and frequency band (stated around pie chart with separations in blue) among all significant predictors (*N* stated in centre) for the indicated task. (**c-d**) Values of absolute (c) or relative (d) predictor variables for DMTS WM (extracted from (a)) in correct (blue) and incorrect (red) trials (left axes). Bars in the background, referenced to by right axis, show absolute values of average predictor weights normalized within each classifier (i.e., connection) coded by their colour. Significance is not indicated as it applies to all shown variables. Blue font, directional connectivity in the *opposite* direction compared to connection-name. Error bars, s.e.m. (**e, f**) Left: Number of variables identified as significant based on ML predictor weight (light green), difference between correct and incorrect trials (medium green), or both (dark green) in dependence on the *P*-value adjustment (y-axes) in the DNMTS (e) and T-maze (f) tasks. Right: Share of predictor variables as depicted in (b) for the *P*-levels indicated in green.

Given the prominence and high predictor weights of *CP* parameters (Fig. 4c-d) - which align with the arising γ-band connectivity immediately after the CP-poke (Fig. 2a-d) - we wondered if the predictability of WM-choice (Fig. 3b) actually relied mainly on identifying a representation of anticipated reward. Therefore, we replicated the decoding analysis for *SP* choices for which animals also expect reward. For the SP, however, average decoding accuracies were – although still above the 50% chance level - considerably smaller, namely 64-68 % and 54-59 % for predictions based on connectivity and local activity, respectively (Supplementary Fig. 6a). While this result shows that the attentional element of the task is more difficult to predict from the available parameters than WM choice, it also demonstrates that the obtained *CP* prediction accuracy was not simply based on representations of motor-action (hole-poking), attention, or reward anticipation. We also repeated the decoding analysis for CP choice with complete omission of all CP parameters. Even though average decoding accuracies decreased significantly for some connections, including the most predictive ones (PFC-dHC, 72%, PFC-MD, 73.9%) - but not for vHC-dHC (72%) - overall accuracies remained far above those obtained from classifiers trained on shuffled labels, and hence above chance level (*P* < 10^-30^ and *P* < 0.002 for classifiers trained on connectivity or local data, respectively; Supplementary Fig. 6b). Overall, these analyses demonstrate that activities along distinct connections and in distinct frequency bands represent encoding (SP), maintenance (delay), and recall (CP) of WM contents in the 5-CSWM task.

### WM-related functional connectivity is highly task-specific in mice

To investigate if such phase-specific connectivity generalizes across tasks, we performed the same analysis for the classifiers predicting performance in the operant DNMTS and the T-maze assays. In both cases, considerably more parameters carried WM-related information than in the 5-CSWM task (Fig. 4a-b). Compared to DMTS WM, T-maze rewarded alternation choice was predicted by a much larger number of predictors, with a prominence of θ- and β-range (as opposed to γ-range) variables, and a considerable proportion of SP-parameters (Fig. 4a-b). This pattern is reflective of the diversity of connectivity parameters that have been associated with T-maze performance in prior studies (Supplementary Table 1), and the relatively detrimental effects of optogenetic manipulations in the SP and delay, as compared to the CP^33, 34, 46^.

Most astonishingly, the combined analysis of all three tasks revealed that *none* of the specific connectivity parameters identified in *one* task bore significant predictive power in the other two tasks, revealing a remarkable task-specificity of such parameters (Fig. 4a). It is, however, possible that this finding is simply caused by a very conservative Bonferroni-adjustment of the *P*-value used as significance threshold (0.05/number of *all* connectivity and activity variables combined; 0.05/1184 for the 5-CSWM, 0.05/888 for the T-maze and 2-CSWM). To test this possibility, we repeated the above analysis while relaxing this adjustment incrementally over four orders of magnitude (Fig. 4e-f). However, the number of identified significant parameters, the relative contribution of different frequency bands, and especially the extreme sparseness of overlap between task-specific predictors changed relatively little (Fig. 4e-f, Supplementary Fig. 7-8). This analysis also revealed that far more connectivity parameters are identified according to their prediction weight than according to their amplitude difference between correct and incorrect trials (Fig. 4e-f). This suggests that the classical approach of correlating behavioural performance with the amplitude of a given metric (Supplementary Table 1) likely misses a sizable proportion of WM-related functional connectivity.

To investigate potential differences or similarities between the time-course of individual parameters during the task, we extracted those spectral connectivity parameters that were predictive across all three assays albeit in different phases: directed dHC➔PFC γ-connectivity and intra-hippocampal δ-range coupling (Fig. 4a). Inspection of the time-course of these parameters over the delay and CP revealed that they behaved rather differently in the individual tasks: dHC➔PFC γ-connectivity showed a transient increase during the delay of all three tasks, but only in the operant tasks a second increase occurred immediately after correct choices (but not after incorrect choices; Fig. 5a-b). Intra-hippocampal δ-coupling even showed a different time course in every task, including a correct choice-specific decrease in the 2-CSWM delay which contrasted sharply with a steady rise during the T-maze delay (Fig. 5a-b). Thus, even within the few predictor variables that are relevant across all tasks, the actual physiological activity relating to the behaviour differed markedly.

**Fig. 5.**
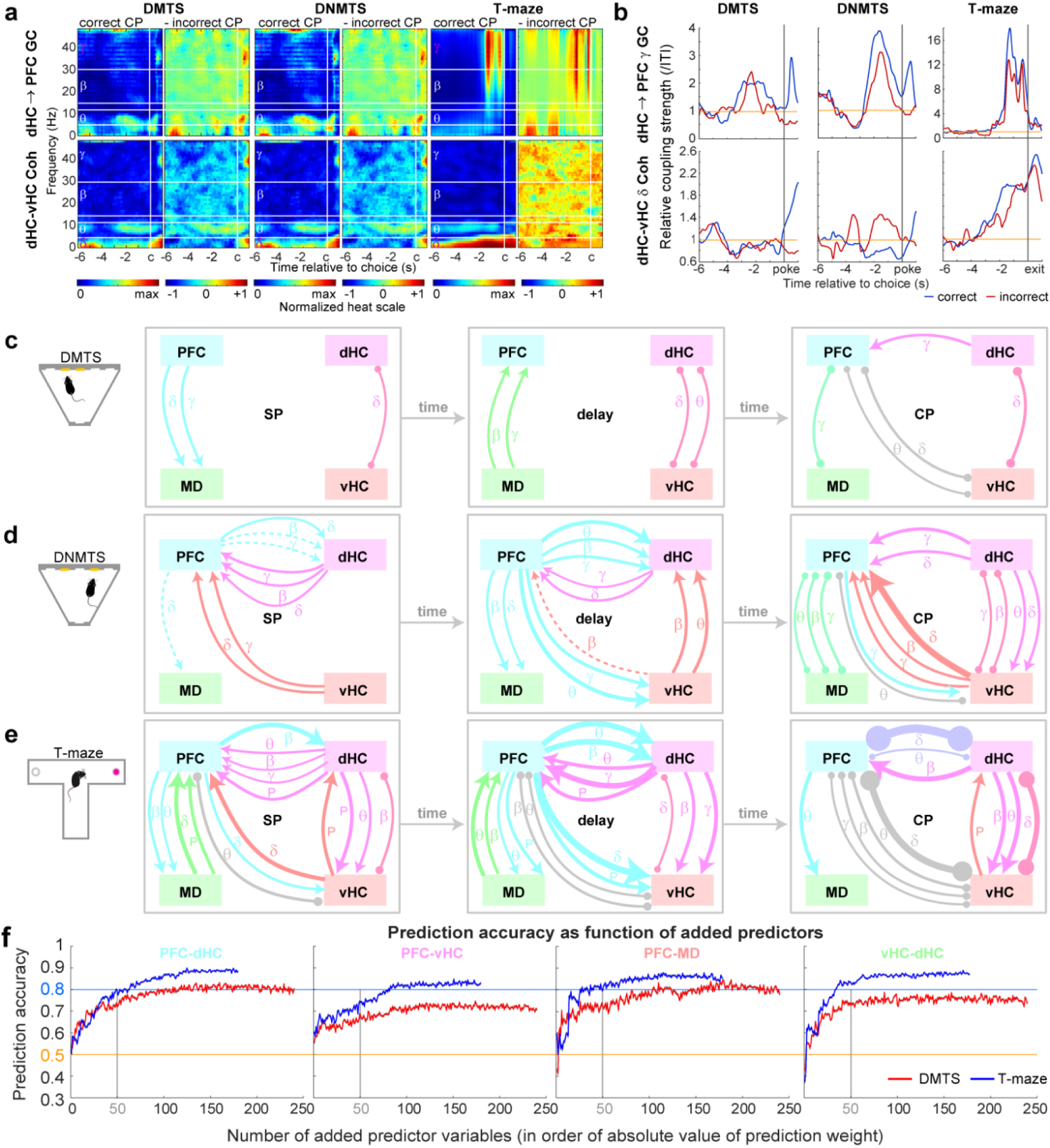
Individual connectivity measures predicting WM choice in all tasks in ice. (**a**) Spectrograms for metrics (stated on the left) that were predictive in all three tasks (albeit in different phases; named at the top) are shown as absolute values and as difference between correct and incorrect CP responses, −6s until +1s around the choice; relevant frequency bands are γ (top) and δ (bottom). (**b**) Average temporal evolution of those metrics in units of their value during the preceding ITI aligned to the choice point in each task (vertical grey line, poke or exit from decision zone) during correct (blue) and incorrect (red) trials. (**c-e**) Depictions of directed connectivity (arrows, derived from GC or PDC) or non-directed coupling (round-ended arcs, coherence or wPLI) during the three phases of each task identified on the left, derived from significant predictor variables shown in Fig. 4a. For connections where there is a directed connectivity metric of the same frequency, a non-directed metric in the same frequency band is not depicted. Line weights indicate the change in connectivity amplitude in the stated phase relative to the preceding ITI (irrespective if the indicated metric was significant as absolute or relative measure). Measures that are significant but where the amplitude in the indicated phase is identical or smaller than in the ITI are shown as dotted lines. P, θ−γ-PAC. (**f**) Average decoding accuracies obtained with classifiers calculated with a reduced number of predictor variables are shown as a function of the number of added predictors, whereby the addition was done in order of normalized prediction weight obtained with all variables (Fig. 3b,c) for DMTS (red) and T-maze (blue) and the named connections. Chance level (50%) and 80% accuracy are indicated by coloured lines. Shaded area, s.e.m.

Given these results, we directly tested the hypothesis that distinct activity patterns underlie the different rodent WM tasks by rendering *task-type* a dependent variable: we trained classifiers to decode which one of the three tasks a subject is currently conducting using connectivity or local activity parameters from correct trials as input. Based on connectivity data, task-type could be decoded with average accuracies of 97-99% when discriminating between the two operant tasks (50% chance level) and with an accuracy >90-95% when discriminating between all three tasks simultaneously (33.3% chance level; Supplementary Fig. 9).

### Common WM-related connectivity patterns shared across rodent tasks

To extract commonalities of connectivity between the three tasks, we aggregated predictive non-directed (coherence, wPLI) and directed (GC, PDC) metrics (extracted from Fig. 4a) and depicted their amplitude increases relative to the preceding ITI for each task phase (Fig. 5c-e). For the T-maze (the only task for which prior reference data exists), this revealed several connectivity patterns associated before with rewarded alternation performance, including vHC-PFC^33^ and dHC-PFC^34^ coupling during encoding and MD➔PFC β-range activity during maintenance across the delay^14, 46^. Importantly, MD-PFC β-range delay activity was also seen in the other two WM tasks, although their directionality differed (MD➔PFC in the DMTS task; PFC➔MD in the DNMTS task; Fig. 5c-e). Likewise, further task-independent connectivity patterns emerged in this analysis: prominent vHC-PFC δ/θ−coupling and vHC-dHC δ−coupling in the CP, and MD-PFC δ coupling in the SP (but with task-dependent directionality; Fig. 5c-e). Some further patterns were shared only by the two DNMTS tasks, e.g., directed PFC-dHC β- and γ-connectivities in the SP and delay (Fig. 5d-e). At the same time, this analysis also confirmed that the vast majority of WM-related connectivity was task-specific, especially when comparing the 5-CSWM DMTS to the other two tasks (Fig. 5c-e).

An important aspect of this analytical approach is that none of these individually highlighted connectivity measures (Fig. 4a, 5c-e) is particularly predictive on its own: When performing decoding analysis with reduced sets of predictor variables – starting with the parameter with the single highest weight and adding variables incrementally – the inclusion of several dozen predictor variables was necessary to achieve maximum decoding accuracy (Fig. 5f).

### Predictive power of local activity in a single area varies by task phase and type

*Local* oscillatory activity in the four analysed regions also allowed considerable prediction accuracy in all three tasks - sometimes even exceeding that obtained from connectivity metrics (Fig. 3b-c). Therefore, to reveal WM-related local activity metrics, we repeated the prior weight-based analysis for the respective variables (power, local PAC). In the 5-CSWM DMTS task only CP parameters, mostly in the β/γ-range, were significantly associated with WM (Fig. 6a). For the two other tasks, in contrast, significant predictors came from all three phases and were somewhat less frequency-specific; the power of dHC-oscillations across all frequency bands and phases constituted the most prominent cluster of choice-predictors in both assays (Fig. 6a). In agreement with the high decoding accuracy obtained with local activity (as opposed to connectivity) in the DNMTS 2-CSWM (Fig. 3b), many more significant local predictor variables were found for this task compared to the other two, irrespective of *P*-value threshold (Fig. 6b). Importantly, however, there was again hardly any overlap between significant predictors from the three tasks. Also, while in DMTS WM all predictive activity parameters had *higher* amplitudes in correct trials compared to incorrect trials, this was not the case for the T-maze, where virtually all predictive hippocampal activity was *lower* in correct trials compared to incorrect trials – only PFC and MD power were higher in correct trials (Fig. 6c-d). Hence, as observed in inter-regional connectivity, local activities related to WM-choice were highly task-specific in multiple respects.

**Fig. 6.**
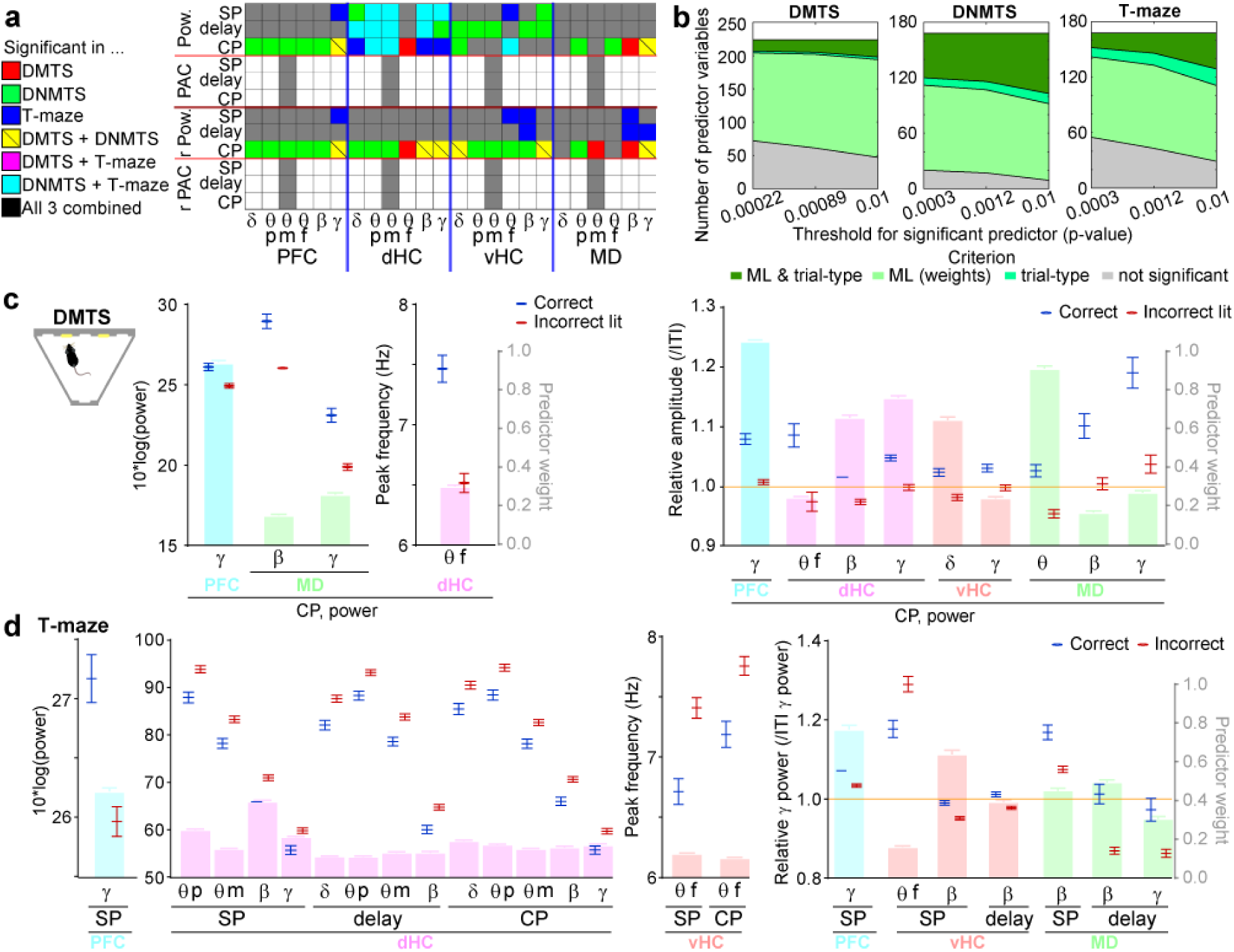
Local activity measures predicting WM choice in all tasks. (**a**) Display and analysis as in Fig. 4a, but for all local activity parameters that contributed to the region-based classifiers shown in Fig. 3b,c. Variables that were significantly associated with WM-performance according to both their prediction weight and differences between correct and incorrect CP (*t*-tests, Bonferroni-adjusted for total number of variables) in any of the three WM tasks are indicated by the corresponding colour, remainder in grey (white squares have no corresponding variable). (**b**) Number of variables identified as significant based on ML predictor weight (light green), difference between correct and incorrect trials (medium green), or both (dark green) in dependence on the *P*-value adjustment (x-axes) in the named tasks. (**d-e**) Values of absolute (left) or relative (right) predictor variables for DMTS (e) or T-maze (f) WM (extracted from (a)) in correct (blue) and incorrect (red) trials (left axes). Bars in the background, referenced to by right axis, show absolute values of average predictor weights normalized within each classifier (i.e., connection) coded by their colour. Significance is not indicated as it applies to all shown variables. Error bars, s.e.m.

### Trial-by-trial decoding of WM-mediated choices from local and long-range neural activity in humans

It remains unclear if highly task-specific and widely distributed WM correlates are only found in rodents or also in human WM. To clarify this question, we used a dataset of intracranial LFP (iEEG) recordings made in 8 human subjects from three sites - PFC, OFC, and MTL – during three types of WM assays whose trials were intermixed within a single test session: identity-related WM (differentiating between identical and novel shapes), spatial WM, and temporal WM (remembering the temporal order of two stimuli; Fig. 7a-b)^20^. For each of the three tasks, we applied the same ML-approach as in mice, generating classifiers that use activity data from four phases (SP, pre-cue- and post-cue delay phases, CP; see task schedule in Fig. 7a) from only a single connection or region at a time.

**Fig. 7.**
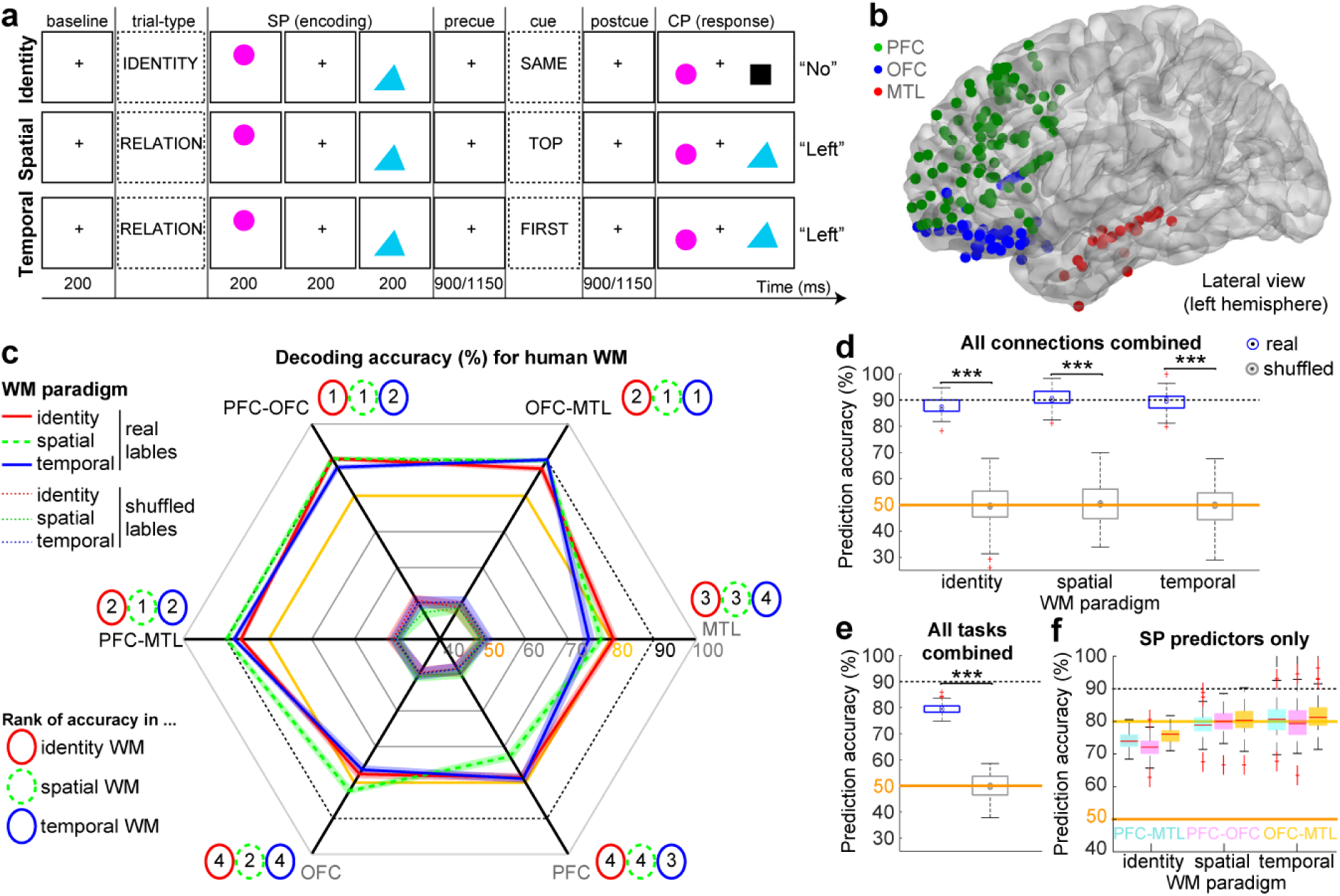
Single-trial based prediction of WM choice in humans. (**a**) Display of the structure of the three human WM tasks, according to ref. 20 (see Methods). (**b**) Placements of electrodes as projected onto the left hemisphere, irrespective of actual hemisphere of each electrode. (**c**) Cross-subject decoding accuracies achieved on average when using connectivity or local activity parameters of the indicated connections (black) or areas (grey), respectively, to predict WM-based *correct* vs. *incorrect* choices in the tasks coded by colour on the left. Accuracies achieved by classifiers trained with randomly shuffled labels are shown as coloured dotted lines; they are consistently lower than accuracies achieved by classifiers trained with real labels in all tasks and connections/regions (*P* < 10^-40^, *t*-tests, not indicated) and assume chance level (50%, orange). Accuracies of 80% (yellow line) and 90% (black dotted line) are indicated to aid comparison. Shaded area represents s.e.m. Numbers in circles colour-coded for the respective paradigm indicate the rank of the average decoding accuracies achieved using data from the respective connection or region. Ranks have been generated from pairwise comparisons with Tukey post-doc tests conducted after significant effects of connection/region in one-way ANOVAs (*P* < 0.0001 in all cases); connections/regions that were not significantly different from each other were assigned the same rank. (**d**) Decoding accuracies achieved when using the classifiers trained on predictors from all connections/regions combined (**e**) Similar analysis as (d) but with trials from all three paradigms inter-mixed. Blue and grey in (d-e) depicts performance of classifiers trained on trials with correct or incorrect (shuffled) labels (stars indicate *t*-test comparisons between them). Error bars, data range without identified outliers which are highlighted in red; boxes, range between 25^th^-75^th^ percentile; dot, median. (**f**) Similar analysis as in (d) but using only SP variables from the connection indicated by colour as predictors. Red lines indicate mean, boxes the 25^th^ and 75^th^ percentile, whiskers indicate data range without outliers, and red crosses indicate outliers. See also Supplementary Fig. 10 for analysis but using only predictors from single task-phases.

Average decoding accuracies for trial-by-trial prediction of WM-choices were higher than those achieved in mice, ranging consistently between 87-90% for predictions based on connectivity and between 72-82% for predictions based on local activity, whereas “predictions” based on shuffled control data remained significantly lower (*P* < 10^-40^, *t*-tests) and were not different from chance level (50%; Fig. 7c). We also trained classifiers on the combined data from all three inter-regional connections and three regions – either separately for each task-type or combining all types of trials indiscriminately. For task-specific classifiers, average decoding accuracies reached 87.6%, 90.8%, and 89.8% for identity-related, spatial, and temporal WM, respectively, i.e., no higher than what could be achieved by connectivity data from the single best connection in each task (Fig. 7d). However, decoding accuracy dropped to 79.4% if task-paradigms were intermixed (Fig. 7e) suggesting that functional connectivity is, at least partially, task-specific. Task-specific prediction accuracies of up to 91% could also be obtained without including CP connectivity measures (Supplementary Fig. 10). Furthermore, in two cases, an average prediction accuracy of up to 81% could even be achieved if using connectivity data from only a single task phase – either the SP in temporal WM or the post-cue delay in spatial WM (Fig. 7f; Supplementary Fig. 10). Strikingly, in both cases, prediction accuracies – and hence information contents - of all three connections were always similar to each other, suggesting a broad presence of WM-related neural substrates across the brain.

### WM-related functional connectivity is highly task-specific and broadly distributed in humans

In order to identify possible correlates of WM in humans, we analysed the prediction weights of the individual connectivity metrics similarly as for the mouse dataset, again extracting WM-related metrics based on the two criteria of prediction weight and a different amplitude of the metric in correct trials compared to incorrect trials (Bonferroni-adjusted *t*-tests). As in mice, WM-related measures (185 out of 1344 connectivity predictor variables) were widely distributed across connections, frequency bands, and metric types. When inspecting the matrix of significant predictors more closely, some regularities emerged (Fig. 8a): First, WM-related activity was highly task-specific with 88% of significantly WM-related connectivity metrics being relevant in only a single paradigm. Only a single metric was predictive in all three paradigms – OFC➔PFC post-cue delay γ-PDC. The principle task-specificity was maintained also with relaxed *P*-value thresholds (Supplementary Fig. 11-12). Second, by far the most – and the most common - predictors emerged in the γ-band, irrespective of significance threshold (Fig. 8a-b, Supplementary Fig. 11, 13-14). The δ-band - in contrast to mice - contributed almost no WM-related variables (only one each in spatial and temporal WM, confined to the OFC-MTL connection). Also, the θ-band bore relatively few WM-related connectivity parameters, and these were mostly relevant for spatial WM and to a lesser extent for identity WM, but hardly for temporal WM. Furthermore, θ-γ-PAC appeared rather relevant (as found in the same data before^20^) in all three types of tasks, especially identity-related WM. Third, *changes* of a metric relative to the ITI before each trial were rarely predictive. Finally, despite the relatively high decoding accuracy achieved for temporal WM (Fig. 7c-d), the number of connectivity metrics related to this WM-type was considerably smaller (18 out of 1344 measures) than for the other two (72 and 95) and there was hardly any overlap between these metrics and those relevant for the other two WM-paradigms (only 3 each, mostly in the γ-band; Fig. 8a-b). In summary, the analysis in humans confirms the high task-specificity, and broad anatomical and frequency-range distribution of WM-related neural activity already seen in mice.

**Fig. 8.**
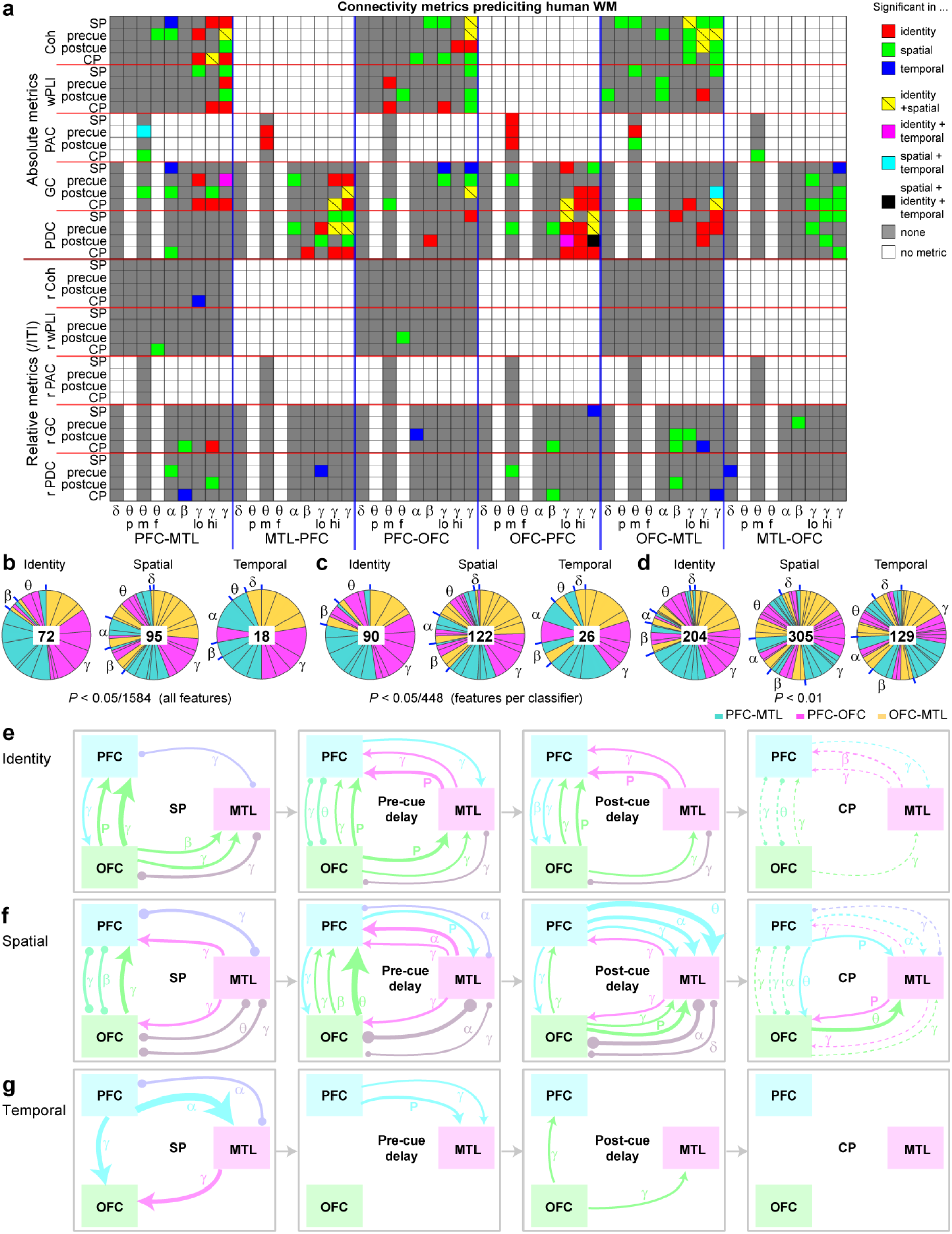
Highly task-specific and broadly distributed correlates of human WM. (**a**) Matrix showing all connectivity predictor variables that contributed to the classifiers shown in (8d) and were significantly associated with WM-performance according to their weight and differences between correct and incorrect CP (see Results) in the paradigms coded by colour (see legend on the right). For θ, mean amplitude (m), peak amplitude (p), and frequency of peak (f) are shown, while for all other variables only the mean amplitude is used. The γ-band contributed three predictors each as this frequency was split into a high- and low-γ-range in addition to using the whole range (30-100 Hz). See Supplementary Fig. 11 for the same analysis with relaxed *P*-value correction and Supplementary Fig. 13-14 for prediction weights of the same variables. (**b-d**) Share of each connection (coded by colour) and frequency band (stated around pie chart with separations in blue) among all significant predictors (*N* stated in centre) for the indicated task. The significance threshold has been Bonferroni-adjusted either by the total number of predictor variables from all connections and regions (1548, corresponding to analysis in panel (a); b), or by the number of connectivity variables for a single connection (448; c), or a standard threshold of 0.01 was chosen (d). (**e-g**) Depictions of directed connectivity (arrows, derived from GC or PDC) or non-directed coupling (round-ended arcs, coherence or wPLI) during the four phases of each task identified on the left, derived from significant predictor variables shown in (a). Line weights indicate the increase of connectivity in the stated phase relative to the preceding ITI. Measures that are significant but *decrease* in the respective phase relative to the ITI are shown as dashed lines. Significant coupling metrics are not depicted if directed measures are represented in both directions. P, θ−γ-PAC.

### Common g-band connectivity across human WM tasks

To scrutinize this conclusion, we searched for commonalities between tasks by aggregating predictive non-directed (coherence, wPLI) and directed (GC, PDC) metrics and the multiple measures within the γ- and θ-bands (extracted from Fig. 8a), and depicted their amplitude change relative to the preceding ITI for each task phase (Fig. 8e-g), as previously done for the mouse dataset (Fig. 5c-e). In this analysis, OFC-PFC γ-coupling during encoding and directed OFC➔PFC/MTL γ-connectivity during the post-cue delay emerged as common patterns present in every task. There were also more commonalities between spatial and identity WM, namely γ-coupling between all three regions that was elevated throughout encoding and delay phases and then decreased below ITI-levels in the CP (Fig. 8e-f). Strikingly in fact, *all* significant predictors from the α−β−γ-range in these tasks showed *elevated* amplitudes during encoding and delay, but *decreased* amplitudes during the CP, compared to their amplitude in the preceding ITI (Fig. 8e-g). Only θ and θ−γ-PAC predictor variables *increased* in amplitude during the CP in spatial WM (Fig. 8f). Finally, even with this aggregated analysis, not a single connectivity pattern that was shared between any two tasks emerged outside the γ-band in any task phase (Fig. 8e-g). This analysis suggests that human WM generally relies on anatomically broad, task phase-specific modulation of γ-connectivity between several brain regions irrespective of task, while the engagement of oscillatory coupling in other frequency bands is task-specific.

## Discussion

Here we demonstrate that WM-related choices can be predicted trial-by-trial in mice and humans using linear decoding of high-dimensional arrays of LFP-based measures of inter-regional connectivity or local activity (the *electome*). The high decoding accuracies of around 90% (compared to a chance level of 50%) achieved in both species are remarkable considering the spatially coarse nature of the extracted neural signal, the short – often sub-second – data traces used to calculate predictors, the intrinsic variability caused by merging data from all analysed individuals with varying electrode placements, and the lack of precise neuronal information as encoded in spike trains of individual neurons^10, 21, 33, 41, 47^. Using the trial-by-trial predictive power of physiological activity as the indicator for its association with WM^42^, this approach enabled a largely unbiased top-down analysis to reveal an unexpectedly rich pattern of frequency-specific connectivity changes during individual phases of distinct WM assays in mice and humans.

The comparative analysis of multiple WM assays - including those that allow control over basic motivational and attentional parameters - provides a unique advantage in that neurophysiological activity patterns which might be truly relevant to WM may be isolated. In this way, we could reveal MD-PFC δ− and β-range coupling during memory encoding and maintenance, respectively, as well as vHC-PFC and vHC-dHC δ/θ-coupling during retrieval as common connectivity patterns across all three rodent tasks - although the vast majority of connectivity proved to be task-specific (Fig. 5c-e). In humans, γ-band connectivity across all analysed connections was commonly linked to WM-choice across tasks, while WM-related connectivity in other frequency bands was mostly task-specific (Fig. 8).

Against a backdrop of widely varying assumptions about which kind of neural connectivity underlies WM (see Introduction), our analysis was initially motivated by the possibility to extract “true” anatomical and frequency-related WM-correlates using the predictor weights generated by the linear classifiers that decode WM-based choices with high accuracy. Our results, however, refute some implicit key assumptions of this endeavour – and, by extension, of many prior investigations of WM-correlates: First, there is no single – or small set of – anatomical regions or connections, types of directional information transmission, or frequency bands that can be regarded as a unique *WM correlate*. Indeed, previously suggested “correlates”, especially in the rodent literature (Supplementary Table 1), could appear as such only because the sum total of connections and measures investigated in each study was small (streetlight effect), as opposed to the 1184 and 1584 metrics analysed here in mice and humans, respectively. In our study, virtually every analysed frequency band, metric, connection and region bore some predictive power regarding WM-mediated choice.

Second, there is no single behavioural WM-task that could be regarded as *representative* of the generic psychological construct termed “working memory” in order to allow the identification of *the* neurophysiological correlate of that construct. The latter is illustrated by the enormous variability in the patterns of predictive connections and metrics across task-paradigms in both species. In other words, a physiological variable that correlates with choice accuracy in the T-maze represents a neurophysiological correlate of T-maze performance, but not necessarily of WM. The same principle applies to our cross-species comparison, as the uncovered candidates for a generic (task-independent) WM correlate originated from different frequency bands in humans (γ) than in rodents (δ−θ−β). A translational implication of these findings is that it is likely impossible to define neurophysiological underpinnings of “working memory” as a uniform psychological construct. However, the existence of certain tasks that represent such a psychological function (i.e., that engages a physiological mechanism that is central to all WM tasks across species) is an implicit key assumption of the *Research Domain Criteria (RDoC)* approach, which envisions to use those representative paradigms in search of WM-enhancing cellular and molecular targets^48^. Our data suggest that the key target variable in the preclinical discovery of WM-enhancing compounds might be the appropriate regulation of γ-range connectivity (given its importance for human WM) rather than behavioural performance in any particular rodent task.

The analysis of the human dataset, in particular, paints a rather different picture of what a correlate of WM could be – at least when searched for in LFP-data. In all three tasks, prediction accuracy calculated from connectivity (as opposed to local activity) metrics was not only very high, but it was also roughly *equal* between the three analysed connections, even though these are anatomically quite distinct. This was the case even with the limited analysis incorporating only SP or post-cue delay connectivity as predictors of spatial WM. These findings may be taken as an indication that WM-related information is extremely broadly distributed, and - rather than specific activity located in a certain connection or region - it is the ability to manipulate information flow across brain regions *as such*, that determines task performance^49, 50^. This model is in line with the ever growing list of brain areas that are implicated in WM, including the superior frontal^51^, anterior cingulate^52, 53^ and sensory cortex^54, 55^, ventral tegmental area^28^, and the nuclei of the midline and anterior thalamus^4, 9, 34^, and the concept that different areas may be involved depending on the strategy used to solve the task^54^. An unexpectedly broad anatomical representation of sensory and behavioural information across the brain has recently been uncovered by decoding activity of individual neurons in multiple cortical areas^56, 57^, and our decoding analysis of LFP-based connectivity in cognition underscores this phenomenon.

In conclusion, our multi-area decoding approach and cross-task cross-species comparative analysis revealed not only a rich functional connectivity supporting WM, exceeding the WM-associated connectivity described before. It also demonstrated an unexpected task- and species-specificity of WM-related neural coupling that raises substantial caution regarding the predictive translational value of each assay and demands to re-think our search for physiological WM-correlates.

## Methods

### Animals and surgery

All experiments were performed in accordance to the German Animal Rights Law (Tierschutzgesetz) 2013 and were approved by the Federal Ethical Review Committee (Regierungsprädsidium Tübingen) of Baden-Württemberg, Germany (licence number TV1399). The rodent cohort, surgery and histology details have been described in our prior publication of the open-field data from the same cohort^37^. Briefly, 12 C57BL/6N wildtype mice, including 8 males, were selected from offsprings of an in-house colony (heterozygous *Gria1*^tm1Rsp^ mice; MGI:2178057)^58^, group-housed in Type II-Long individually ventilated cages (Greenline, Tecniplast, G), enriched with sawdust, sizzle-nest^TM^, and cardboard houses (Datesand, UK), and subjected to a 13 h light / 11 h dark cycle. The mice were implanted with electrodes at ca. 9 months of age, after 2 months of training in the DMTS-WM task and prior habituation training (see below). Single polyimide-insulated tungsten wires of 50 µm diameter (WireTronic Inc., CA, US) were implanted, with reference to Bregma (in mm), into the PFC (AP +1.8-1.9, ML 0.3-0.35; 1.8-1.9 below pia), MD (AP −1.2, ML 0.3, 2.7 below pia), dHC (AP −1.9-2.0, ML 1.5, 1.4 below pia), and vHC (AP −3.1-3.2, ML 2.9-3.0, 3.4 mm for single and 3.8-3.9 mm for dual electrodes below pia). In most mice, dual electrodes were used for PFC and vHC, whereby the second electrode was placed about 0.5 mm higher than the stated distance from pia. Both hemispheres were implanted at roughly equal proportion. Stainless steel screws (1.2 mm diameter, Precision Technologies, UK) were implanted in the contralateral hemisphere ca. 1 mm from the midline above the cerebellum (AP −5.5) for ground and above the anterior frontal cortex (AP +4.0) for additional reference (used for the analysis in the open-field test, but not for the present analysis), and where connected with a 120 µm PTFE-insulated stainless steel wire (Advent Research Materials Ltd., UK; Fig. 1b). All electrode wires were connected to pins in a dual-row 6-pin or 8-pin connector (Mill-Max, UK). Electrode placements were determined *post-mortem* from electrolytic lesions made under terminal ketamine/medetomidine anaesthesia followed by perfusion-fixation. Misplaced electrodes were excluded, leaving the following number of used electrodes per region; PFC: 12; dHC: 7; vHC: 9; MD: 4. The resulting number of contributing connections were PFC-dHC: 7; PFC-vHC: 9; vHC-dHC: 6; MD-PFC: 4.

### Operant DMTS 5-CSWM task

The principal schedule of the task was as previously described^45^ and emulates the *combined attention and memory* (CAM) task previously developed in rats^59^. In the present study, the task was conducted in custom-designed pyControl-based operant boxes that were optimized for both simultaneous electrophysiological recordings and the acquisition of the 5-CSWM task (operant box design-files and task-scripts implementing all task paradigms used in this study available from https://github.com/KaetzelLab)44. The latter was done by using a 5-choice poke-wall with a larger distance between the holes (3 cm edge-to-edge; 4.5 cm centre-to-centre), which we chose because of the long learning time in our standard commercial boxes used before^45^. Briefly, each trial of the task is divided into a sample phase (SP), a delay-phase (delay) - which is further sub-divided into a pre-delay and a post-delay by the time point of reward collection - the choice phase (CP), and the inter-trial interval (ITI). In the default state, the task is conducted in the dark (house-light off) with the only illumination deriving from the poke- and receptacle holes in certain task phases. The SP is identical to that of the 5-choice-serial-reaction-time task (5-CSRTT), except that premature responses are not punished: one of the 5 holes in the 5-choice wall is illuminated for a certain stimulus duration (SP-SD) and mice need to poke into that hole within the limited hole time (SP-SD plus 1 s) in order to obtain a small reward (20 μl or, in stages 2 and onwards, 10 μl strawberry milk) at the receptacle at the opposite end of the wall during the pre-delay time. The reward-collection (receptacle exit) starts the post-delay (2 s in all cases) after which the originally presented hole and one randomly assigned other hole is illuminated (thereby starting the CP) for a certain stimulus duration (CP-SD, 5 s for all protocols shown in Fig. 1). Mice have to poke into the same hole as in the SP, realizing a DMTS-rule, in order to obtain a large reward (60 μl). After a 5 s ITI a new trial starts with the SP. Incorrect pokes or omissions in the SP or CP lead to a 5 s time-out period (house-light illuminated; reward omitted) and the start of a new trial after an ITI of 5 s. Before training, mice were habituated to the operant box, to consuming the strawberry milk (Müller®, Germany) reward from the receptacle, and to poking into the 5-choice wall to obtain a reward (acquisition of the basic operant cycle).

Subsequently, training was conducted through multiple stages across which the task became incrementally more difficult due to a shortening of SDs and an increase in the number of CP-stimulus configurations (see Supplementary Table 5). During pre-surgery training – in order to compare performance between groups – no performance-based staging was applied, but instead all mice were trained on the stage 1 for the same number of 21 days, and then transitioned through the remaining three training stages (2-4) with 2-3 d of training per stage. Parameters defining the stages are found in Supplementary Table 5. Sessions lasted 30 min throughout, and were conducted on 5 d/week. Training was continued ∼4-5 weeks after surgery on the baseline stage (4) for 5 weeks to allow the mice to approach asymptotic performance. Subsequently, mice were trained further for 3 weeks in a tethered mode, with the headstage (see below) mounted on their heads, in a baseline protocol with shorter SD in SP (10 s) and CP (5 s; stage 5) to increase the number of obtained trials and better standardize the encoding time. Subsequently, mice were taken through three series of challenge protocols with intermittent training on the baseline stage 5. The same challenge was conducted on 2-3 consecutive days in order to obtain sufficient trials for later analysis. The three series were *(a)* pure delay challenges, where the pre-delay was extended from 0 to 5 and 10 s, *(b)* a distraction challenge with an illumination of the house-light for 0.5 s starting randomly timed between 0.8-1.3 s of the 2 s post-delay phase, and *(c)* a combined attention (SP) and working memory challenge with an SP-SD of 1 s (instead of the 2 s of the specific baseline stage 6 of this challenge) and a pre-delay of 5 s, which was preceded by a sole attention challenge (1 s SP-SD, 0 s pre-delay). Most of the analysis shown uses the data from the final (combined) challenge, although decoding analysis was also conducted for the other challenges in order to replicate the analysis and, additionally, to assess the capacity of cross-prediction between classifiers from entirely different challenge-conditions (Supplementary Fig. 5). The post-delay remained 2 s, starting with the exit from the reward receptacle and spent in darkness, throughout all challenge and baseline protocols.

### Operant DNMTS 2-CSWM task

The operant DNMTS task followed the same principal trial-schedule as the 5-CSWM task except for two modifications: the implementation of a *non*-match-to-position rule (i.e. animals are rewarded for choosing the illuminated CP poke-hole that is not the one, that they poked in the SP) and a simplified set of choice options using always only holes 2 and 4 of the 5-choice wall (making it similar to a task developed by Goto & Ito^60^). Mice were trained in this task only after the T-maze (see below) in order to ease the switching from the prior, opposite task-rule. All mice were trained for 30 training sessions on the 2-choice DNMTS task, then were tested in two delay challenges, in which the pre-delay was extended to 5 and then 10 s (2 d each). A subset was then trained in a 5-choice version of the same task (i.e. the DNMTS-version of the 5-CSWM task) for 12 d *without* and for 3 d *with* headstage mounted, but performance in many mice was not sufficient, and hence mice were returned to the 2-choice version and trained for a further 5 d with mounted headstage. The other subset moved directly to 5 d of training with mounted headstage. Then, recordings were conducted on two days with baseline training and two days each for two delay challenges, in which the pre-delay was extended to 5 and then 10 s. Given the relatively low performance in the delay challenges, the data from the two baseline days was used for further analysis of electrophysiology data. In all protocols, the SP-SD was 8 s, the CP-SD was 5 s, the limited hold time exceeded the SD by 1 s, ITI and time-out were each 5 s, the post-reward delay was 2 s (spent in darkness), the SP reward was 10 μl, and the CP reward 60 μl.

### Non-matching-to-position T-maze rewarded alternation SWM task

Rewarded alternation-based SWM was tested in a T-Maze with transparent walls (10 cm high), intransparent floor (red PVC), and food wells (made from white Teflon®) placed in the end of all three arms. Sliding doors could be used to block off either of the two choice arms in the SP and the initial 10cm partition (containing the food-well) of the start arm during the delay and ITI. Before testing mice were habituated to the surroundings and to the condensed milk reward (10% Ja!-Kondensmilch®, Germany, diluted 1:1 with drinking water) at first cage-wise and later independently. The task schedule was identical to what we previously described^26^, involving ten trials per day each consisting of an SP and a CP. During the SP, mice were placed at the beginning of the start arm facing the experimenter, and were left to run into the pseudo-randomly assigned goal arm that was not blocked to obtain a reward. Subsequently, mice returned to the end of the start arm to obtain another, small reward while being enclosed for a 5 s delay period, during which the door from the previously blocked goal arm was removed. Once the delay had passed, the CP started in which the mouse was allowed to choose between one of the goal arms of which the previously unvisited one was rewarded (correct choice), while the other one was not (incorrect choice). Once the CP was completed, mice were motivated to return to the start arm to obtain another small reward there while being enclosed for an ITI of 20 s before the next trial commenced. 10 consecutive trials were conducted per day; testing was conducted over 8 d with a delay of 5 s and for another 4 d with a delay of 30 s – all 12 sessions were conducted with simultaneous recording of LFP-activity, i.e. a headstage was mounted and tethered through an SPI-cable. In each session, a new sequence of open arms in the SP was used whereby half of the 10 trials were always assigned to the left arm, and the sample arm was not the same for more than three consecutive trials.

### Acquisition and pre-processing of mouse data

Prior to testing, a 32-channel RHD2132 headstage (Intan Technologies, CA, US) was plugged into the implanted connector via a custom-built adaptor that interfaced a 36-pin Omnetics connector (A79022-001, MSA components, G) with another 6-pin or 8-pin Mill-Max connector. The headstage was connected to an *Open-Ephys* acquisition board (https://open-ephys.org, US; obtained through the Open-EPhys store at Champalimaud, Portugal) via two light-weight flexible SPI-cables (Intan Technologies), sometimes daisy-chained through a custom-connected miniature slip-ring (Adafruit, NY, US). The adaptor was wired so that all signals were referenced to the ground-signal obtained from above the contralateral cerebellum. Data were amplified and digitized, sampled at 20 kHz and band-pass filtered at 0.1 – 250 Hz for all subsequent analysis of LFP signals. In operant tasks, all individual task- and behavioural events were recorded by pyControl. Additionally, all events relevant to time-locked electrophysiological analysis of WM task-phases and choices (e.g., correct SP and CP responses) were encoded as patterns of transistor-transistor logic (TTL) signals by pyControl and recorded as time-stamps with the electrophysiological data by the *Open-EPhys* acquisition software using the 8 analogue inputs of the acquisition board and a dedicated BNC-HDMI interface board (Open-EPhys). For the T-maze task, ANY-maze (Stoelting) was used to track the position of the animal in the different subdivisions of the maze, and this positional information was encoded in patterns of TTL-signals recorded via an AMi-interface board and a BNC-HDMI interface board as time-stamps with the electrophysiological signals in the *Open-EPhys* acquisition software.

### Human intracranial electrophysiology data during WM

A publicly available dataset of multi-site intracranial recordings during three WM tasks ^20^ was downloaded from http://dx.doi.org/10.6080/K0VX0DQD. It includes data from 10 adult human subjects (mean ± SD [range]: 37 ± 13 [22–69] y; 7 males) who were implanted with intracranial electrodes to identify epileptic foci for surgical resection. Electrode placements were in the medial temporal lobe (MTL, i.e., CA1; CA3/dentate gyrus; subiculum; or parahippocampal, perirhinal, or entorhinal area), lateral PFC (inferior, middle, or superior frontal area), and OFC (orbitofrontal, frontal polar, or medial prefrontal area). Only subjects with electrodes localized in all three regions were included in the present analyses (*N* = 8). Details of behavioural testing, data acquisition, and pre-processing have been described previously^20^. In brief, three different WM tasks were conducted, whereby subjects had to either identify a previously indicated object identity, location, or temporal order of two visual stimuli. Trials from all these three tasks were pseudo-randomly mixed from trial to trial in a single test session. Each trial started with a 1 s pre-trial fixation interval, after which a screen indicated whether in the respective trial would be tested for object identity or spatiotemporal position (800 ms). Subsequently, the SP started in which two shapes were presented subsequently for 200 ms each, separated by 200 ms. After a subsequent pre-cue delay (900 or 1150 ms, varied pseudo-randomly), a cue appeared for 800 ms that specified which of the two shapes would need to be identified in the later CP according to a rule of identity (same/different), spatial location (top/bottom) or temporal order (first/second). After a post-cue delay (900 or 1150 ms, varied pseudo-randomly), the CP started as two shapes were presented, of which the participant had to choose the one that was correct according to the prior cue. All trials from all patients were merged for subsequent analyses, just as was done for the mouse WM data. The fully pre-processed data (as described^20^) was used for the current analysis.

### Data analysis

All signal analyses were done in MatLab (MathWorks). Mouse electrophysiology data were exported to MatLab and, for all LFP analyses, down-sampled to 1 kHz and analysed with custom-written scripts. To reduce low frequency drift, signals were first detrended using the *locdetrend* function of the Chronux signal processing toolbox (http://chronux.org/) with 1 s of data and a sliding window of 0.5 s. Trials were excluded from further analyses if the amplitude exceeded the 5^th^ standard deviation within each channel for more than 10% of the trial duration. In mice, PFC and vHC were recorded with dual electrodes. In case both electrodes were located in the intended target area (PrL/Cg1 for PFC and fissure for vHC), as inferred from lesion sites, all metrics were calculated for both electrodes and the result was averaged to obtain the final value. For the human dataset, the number of electrodes per site varied between 1 and 28 per area. Therefore, for connectivity measures, each single metric was calculated for every possible inter-regional pair of electrodes and the resulting value was averaged across all combinations of a single connections. Analogously, for local activity measures, each single metric was calculated for every electrode and the resulting value was averaged across all electrodes of a given area for each subject.

#### Power and non-directional synchrony

Power and coherence spectra were calculated with Chronux routines implemented in the Chronux toolbox using the multi-taper method^61^. Power values were expressed as 10*log_10_ values for all analyses and the range of frequencies was set from 0.1 to 48 Hz. A time-bandwidth product of 9 and 17 tapers were used to calculate power and coherence during defined time-periods during ITI, sample phase, delay (if applicable) and choice phase. To address the issue of volume conduction, we calculated the weighted phase lag index (wPLI)^62^ using routines implemented in the FieldTrip toolbox^63^. The original trials were further divided into 99% overlapping “pseudo trials” with a length of 600 ms and padded to the next power of two. The complex cross-spectrum was computed using a Hann taper with a spectral smoothing of 2 Hz. Time-frequency spectral analyses were performed with routines from the FieldTrip toolbox using Morlet wavelets with a width of 3 cycles steps of 10 ms^63^. Time periods before and after the time frame of interest were padded with real data to avoid artifacts of too long wavelets at low frequencies.

#### Directional Synchrony

Non-parametric Granger Causality (npGC)^64^ and partial directed coherence (PDC)^65^ were calculated using the FieldTrip toolbox^63^. The original trials were further divided into 50% overlapping “pseudo trials” with a length of 1s and padded to the next power of two, differing from power and non-directional synchrony measures because 99% overlap did not provide substantially different results but came with a much higher computational effort. The complex cross-spectrum was computed using a Hann taper with a spectral smoothing of 2 Hz. The noise covariance matrix and transfer function were obtained by applying Wilson’s spectral matrix factorization to complex Fourier-spectra. This non-parametric approach was shown to be better at capturing all spectral features, less error prone because no model order had to be chosen, computationally faster than applying autoregressive modelling^66, 67^, and to deliver virtually same results when used on our LFP data^37^. Time-frequency representations of npGC and PDC were obtained by Morlet wavelets using the same configurations as described above.

#### Phase-amplitude coupling (PAC)

Cross-frequency coupling (CFC)^68^ was assessed using the measure of phase-amplitude coupling (PAC), the statistical relationship between the phase of a low-frequency and the amplitude of a high-frequency component, in a cross-regional analysis^69, 70^. Time series data were first band-pass filtered in the desired frequency ranges, followed by a Hilbert transform using the MatLab function *hilbert* to calculate the real and imaginary parts of the signal to obtain the instantaneous amplitude and phase. For each trial, intra- and inter-regional PAC were determined by calculating the modulation of γ-amplitude by θ-phase using the phase-locking technique proposed by Voytek et al. with routines described in^71^.

### Supervised Machine Learning

To validate the calculated measures of neural connectivity and to identify predictive variables we employed supervised machine learning algorithms. Spectrally resolved parameters (e.g., γ-coherence, γ-power) from each inter-regional connection (e.g., dHC-PFC) and local brain region (e.g., dHC) were analysed separately using different classifiers. For classification, we used the absolute parameter values as well as the ratio of each parameter relative to the preceding ITI. In mice, 240 connectivity metrics per connection and 56 local activity metrics per region were used in the 5-CSWM DMTS task (1184 for all connections and regions combined; 296 per task-phase), and 180 connectivity metrics and 42 local activity metrics were used on the operant 2-CSWM DNMTS task and the T-maze (888 combined). The difference originates from the usage of 4 task phases, including pre- and post-delay phases, in the 5-CSWM task, but only 3 phases (one delay phase) in the two DNMTS tasks, which was the post-delay in the 2-CSWM task. For all predictor matrices, the pre- and post-delay phases were combined to allow uniform comparisons between all three tasks. In humans, 448 connectivity metrics per connection and 80 local activity metrics were generated for the analysis shown in Fig. 7, using all 4 task phases including SP, pre-cue delay, post-cue delay, and CP (classifiers omitting the CP were also calculated, see Supplementary Fig. 10). Note that, in humans more predictors arise because frequency bands have been determined additionally in the alpha-band (8-12 Hz), and in a higher γ-band (50-100 Hz) in addition to the low γ-band (30-49 Hz) used in rodent analysis. We also included the combined band (30-100 Hz) as separate predictor. To ensure a sufficient number of trials for classification and the general validity of the identified predicting variables, we merged data from all subjects of one group, i.e., from all mice provided correct electrode placement (see Supplementary Table 6) or from the subset of human subjects with coverage of all three regions, respectively.

Since rodent and human subjects performed proficiently above chance level resulting in more correct than incorrect trials^20^, we used a *synthetic minority over-sampling technique* (SMOTE) to construct a balanced dataset with five nearest neighbours to consider^72^. All electrophysiological predictor variables of a classifier were normalized between 0 and 1, setting the maximum empirical value for each metric to 1. We used 90% of the data as a training set and the remaining 10% for testing. Allocation to the training and test set was done randomly and repeated 100 times to obtain a mean and its variance for the achieved decoding accuracy the predictor weights. To identify the classification algorithm which fits our data best we assessed the 25 most used classifiers implemented in MATLAB (Supplementary Fig. 4). Focussing on easy interpretability and high predictive accuracy we chose random subspace ensembles on a linear discriminant analysis (LDA) template (subspace discriminant classifier) for our further analysis, which achieved the highest prediction accuracies compared to all other tested linear classifiers. Briefly, LDA aims to identify a hyperplane that maximizes the mean distance between the mean of the two classes while minimizing variance between them. Since the sample size of our data was relatively small compared to the number of features, we used the *random subspace method,* which is a valid approach to resolve this issue and has been shown to be superior to single classification algorithms^73, 74^. It operates by creating a classifier ensemble where each classifier is trained with a reduced, randomly sampled number of input features, e.g., they are projected into a new subspace which leads to a relative increase of the number of samples. The number of features to sample in each classifier and the number of learning cycles were set to half of the total number of features and 30, respectively. The coefficient magnitudes of each feature obtained by each subspace LDA classifier were averaged across learning cycles to get a solid quantification of its predictive value. As measures of classification performance, we used the prediction accuracy, the AUC of the receiver operating characteristic (ROC) and the F1-score which is defined as follows: F1 = (2*class1precision * class1recall / (class2precision + class1recall)). Precision is defined as the True positives / True Positive + False Positive for class 1 and as the True negatives / True Negative + False Negative for class 2. Recall is defined as True Positive / True Positive + False Negative which is equivalent to the sensitivity for class1 and specificity for class2. Since it has been shown that the theoretical chance level of 50% should not be expected and it is favourable to obtain an empirical chance level, we randomly shuffled the data labels (e.g. correct / incorrect) and repeated the analysis described above to create an empiric null distribution^75^.

### Statistical analysis

Behavioural training and challenge data were analysed with repeated-measures ANOVA and pairwise Sidak post-hoc tests for simple main effects. To determine the importance of individual connectivity or activity measures (predictors), we used a two-step procedure: First, pre-classification, we performed a paired *t*-test for each feature comparing its value in correct vs. incorrect trials. Second, post-classification, we used the magnitude of the weight of each predictor as delivered by the subspace discriminant classifiers to perform a paired *t*-test between the obtained magnitudes and the magnitudes from the shuffled dataset. Features were only recognized as significantly important if both criteria were met. *P*-values were Bonferroni-corrected within-species for the total number of used features across all classifiers calculated in mice (1184; *P* < 0.05/1184) and humans (1584; *P* < 0.05/1584) in the most conservative analysis; additional analysis was conducted to evaluate the dependency of the number of obtained significant predictors with less stringent adjustments.

### Data and protocol availability

All source data for behavioural performance in mice can be obtained from the corresponding author upon reasonable request. All electrophysiological data from mice will be made publicly available at https://gin.g-node.org/KaetzelLab76. Human data are available at http://dx.doi.org/10.6080/K0VX0DQD. All MATLAB analysis scripts for spectral analysis are publicly available on GitHub (https://github.com/KaetzelLab/LFP_analysis)77. Design files of custom-made operant boxes (https://github.com/KaetzelLab/Operant-Box-Design-Files) and task-files for operant WM tasks (https://github.com/KaetzelLab/Operant-Box-Code) are available on GitHub.

## Acknowledgments

We thank Stefanie Schulz for assistance with histology and Dr. Rolf Sprengel (Heidelberg) for providing the *Gria1*^tm1Rsp^ mouse line. <u>Funding</u>: This work was funded by the Else-Kroener-Fresenius/German-Scholars-Organization Programme for excellent medical scientists from abroad (GSO/EKFS 12; to D.K.), the Juniorprofessorship programme of Baden-Württemberg (to D.K.), the DFG (KA 4594/2-1; to D.K.), the Brain and Behaviour Research Foundation (NARSAD Young Investigator Award 22616 to D.K.), the Alfred-Krupp Foundation (to B.L.), and the National Institute of Neurological Disorders and Stroke (K99NS115918 to E.L.J.).

## Author Contributions

SKTK and DS conducted all mouse experiments. DS analysed mouse and human data, with initial support from AMB and BFG. DS and DK designed the study based on prior input from AMB, BFG, and DMB. BL provided essential resources. ELJ provided the human data. SKTK, TA, and DK developed the pyControl operant box enabling simultaneous recording during operant behavioural testing. DS and DK wrote the manuscript with advice from all authors.

