## Supplementary Information for "Highly task-specific and distributed neural connectivity in working memory revealed by single-trial decoding in mice and humans"

### Supplementary Figures

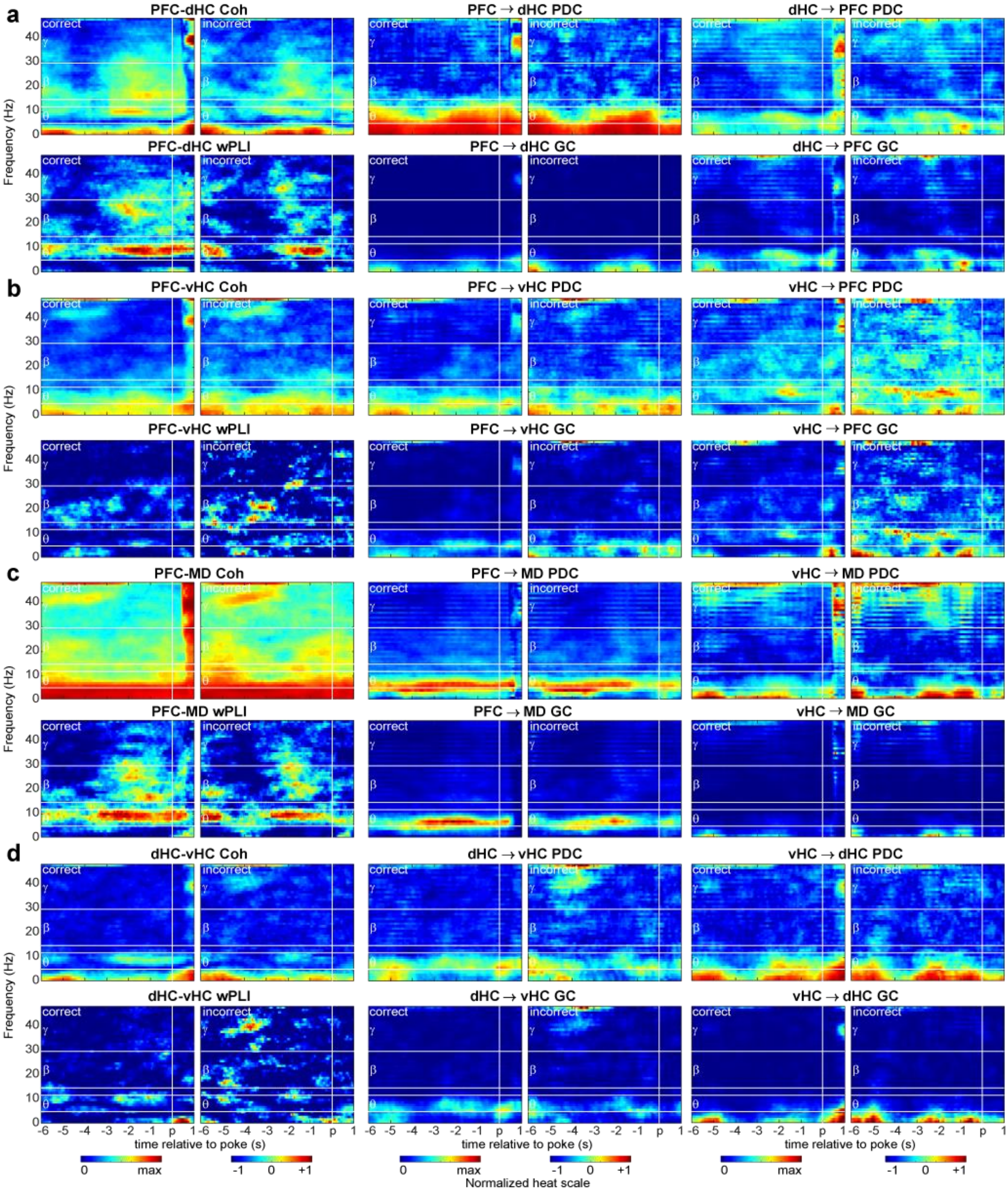


**Supplementary Figure 1 | Spectrograms of connectivity during the 5-CSWM delay and CP (1s SD+5s delay challenge). (a-d)** Spectrograms displayed from 6s before to 1s after a CP poke for the connections and metrics named in subpanel titles. Left and right graphs of each subpanel correspond to connectivity during correct and incorrect responses, respectively. White vertical lines indicate time of CP poke, while white horizontal lines differentiate between frequency ranges named on the left.

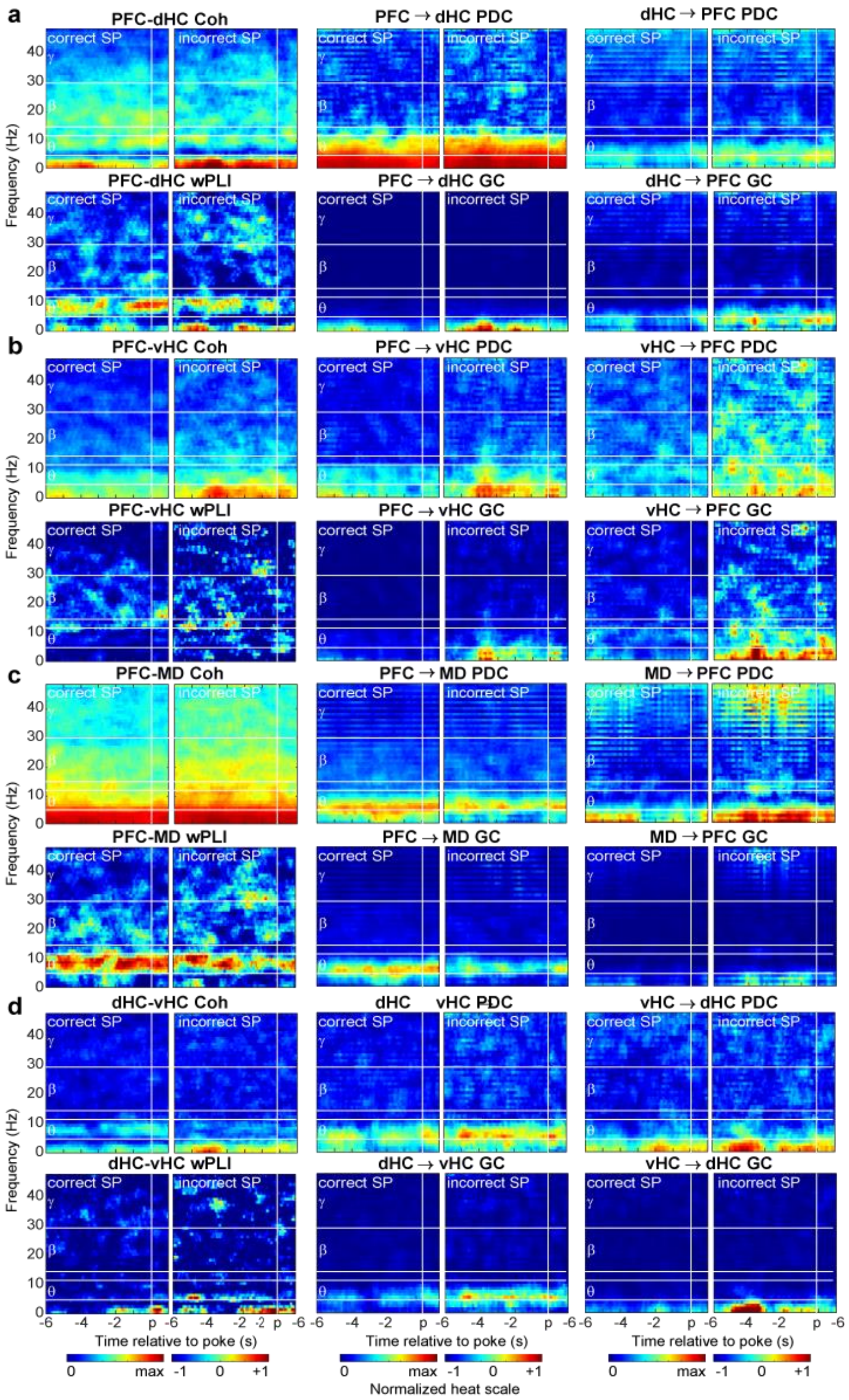


**Supplementary Figure 2 | Spectrograms of connectivity during the 5-CSWM SP (1s SD+5s delay challenge).** Same display as in Supplementary Fig. 1, but for SP. Analogously, left and right graphs of each subpanel correspond to connectivity during correct and incorrect SP responses, respectively. White vertical lines indicate SP poke.

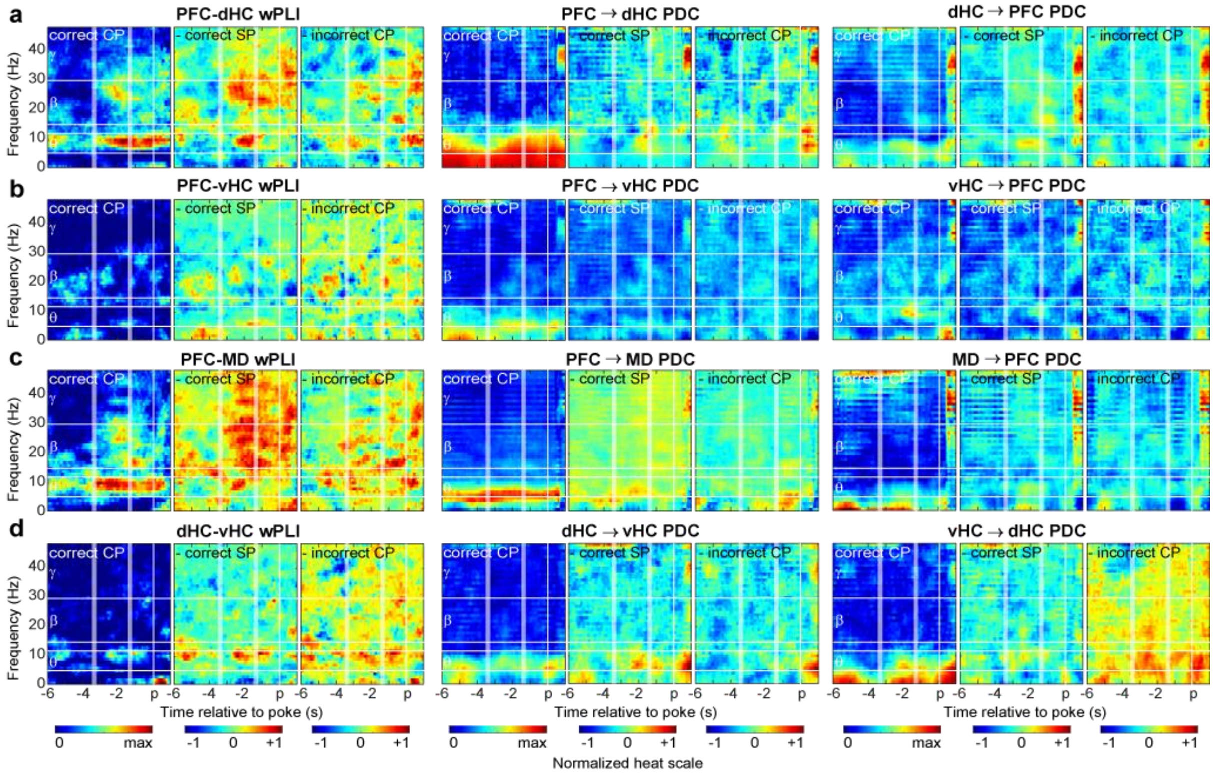


**Supplementary Figure 3 | Spectrograms of differential connectivity during the 5-CSWM delay and CP (1s SD+5s delay challenge). (a-d)** Same display as in main Fig. 2a-d, but for wPLI and PDC. Spectrograms depicting min-max normalized coherence and GC for the connections stated above each triplet panel for the delay and CP of the 5-CSWM task, temporally aligned to the choice poke entry (p, white vertical lines; showing 6 s before until 1 s after the poke); the start and end of the post-delay shown by white stripes corresponding to mean±SD as determined by CP response latency. Each triplet shows the absolute value (left), the difference between the former and either the prior correct SP (controlling for representations of attention and reward, middle), or incorrect CPs (showing differences of WM engagement and subtracting motor representations of poking; right). Horizontal white lines show borders between analysed frequency bands, stated on the left.

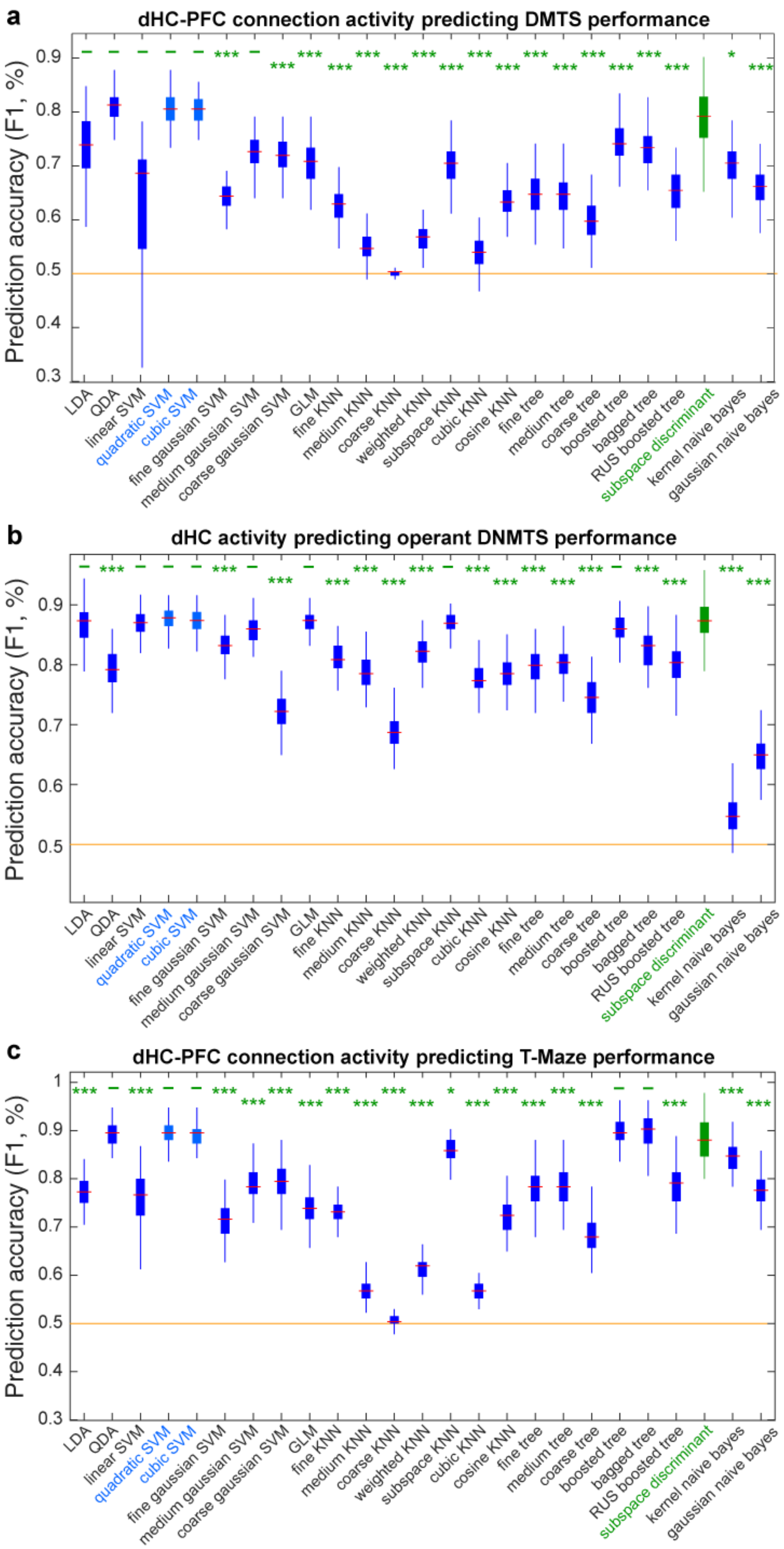


**Supplementary Figure 4 | Decoding performance of different types of classifiers.** (**a-c**) Decoding accuracy displayed as the F1-score for the connections (a, c) or brain region activity (b) which yielded the highest predictive power in its respective task (see main Fig. 3), as named above each panel. The most commonly used classifiers implemented in MATLAB were trained and tested with the same data, and prediction accuracies were compared. Yellow horizontal line depicts chance level. Red lines indicate mean, coloured boxes the 25$^{th}$ and 75$^{th}$ percentile and whiskers indicate data range across the 100 classifiers computed for each type and connection. The subspace discriminant analysis (green) was superior to all tested linear classifier types and therefore used for all analyses in this study. Only two of the tested non-linear classifier types, named in light blue font, yielded equivalent accuracies across all three comparisons (a-c), but non-linear classifiers do not allow to extract and compare predictor weights for individual metrics. Classifier performance was compared using pairwise Tukey post-hoc tests after a significant main effect of classifier type in a one-way ANOVA ($P < 0.0001$). Results of Tukey tests comparing each classifier type to the subspace discriminant type are indicated by green dashes ($P > 0.05$, n.s.) or stars, * $P < 0.05$, *** $P < 0.001$.

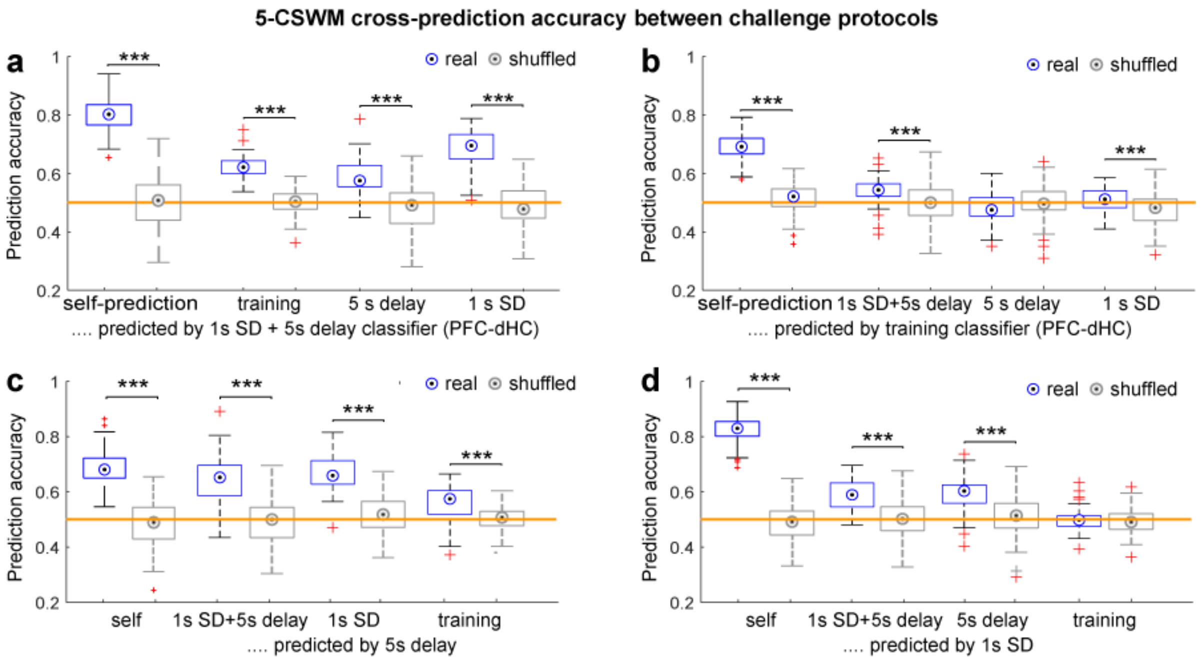


**Supplementary Figure 5 | Cross-prediction accuracy for dHC-PFC classifiers trained on different challenges.** (**a-d**) Decoding accuracies achieved when using the classifiers trained on dHC-PFC connectivity data from the combined challenge (**a**), the training (**b**), the 5 s delay WM challenge (**c**), or the 1 s SP-SD attention challenge (**d**) on data from the other respective 5-CSWM protocols named on x-axes; blue and grey depicts performance of classifiers trained on trials with correct or incorrect (shuffled) labels (stars indicate *t*-test comparisons between them). Error bars, data range without identified outliers which are highlighted in red; boxes, range between 25$^{th}$-75$^{th}$ percentile; dot, median. *** $P < 0.001$

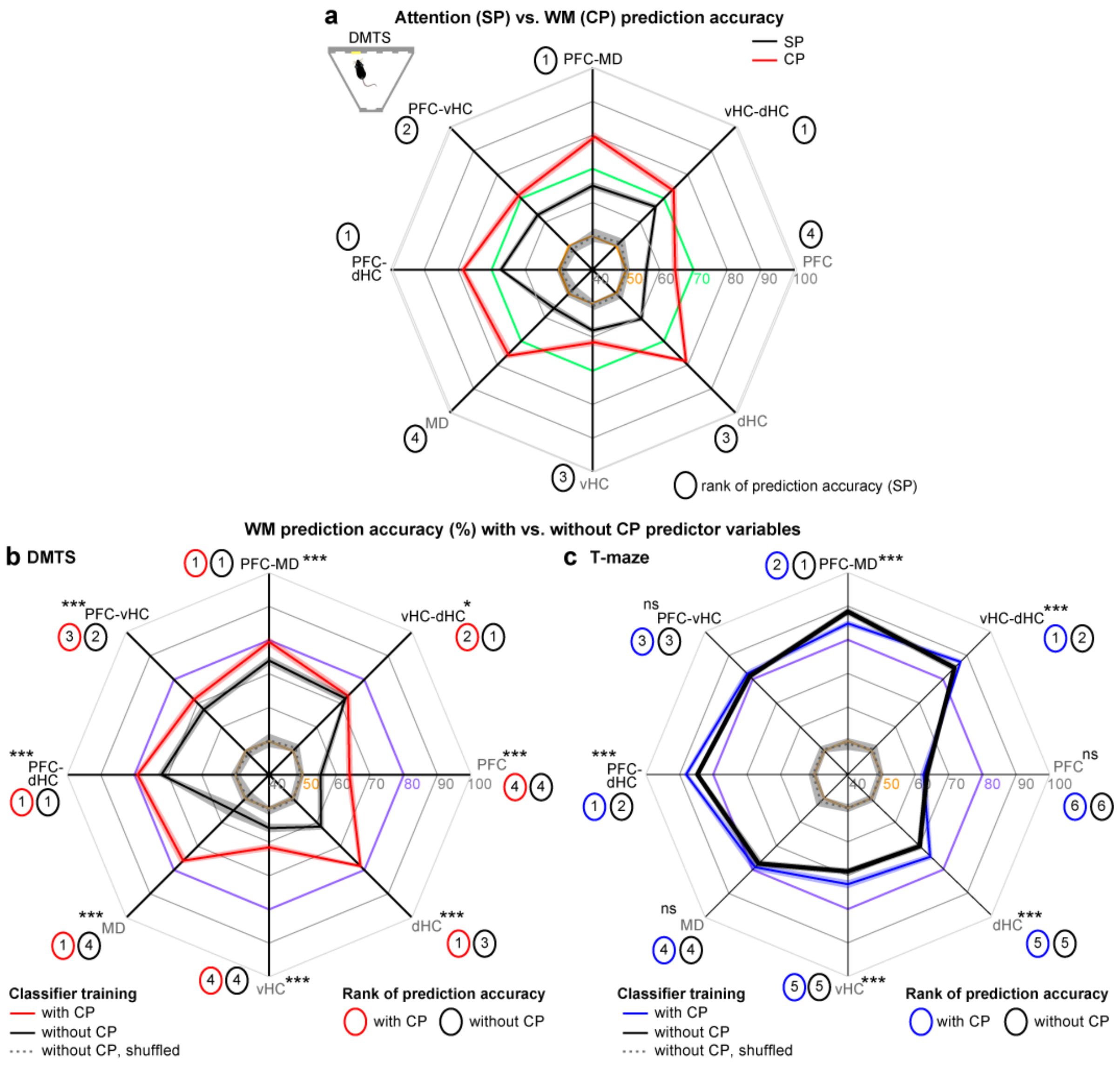

**Supplementary Figure 6 | Prediction accuracies in the 5-CSWM SP and without CP predictors.** (**a**) Cross-subject decoding accuracies for predicting of correct vs incorrect *SP*-choices (as a measure of sustained attention) from SP connectivity or local activity parameters of the indicated connections or areas in the 5-CSWM (black line; combined 1 s SP-SD, 5+2 s delay challenge, 2 sessions). A time window that ends 1 s after the SP-poke was used to retain equivalence to CP-based predictors contributing to WM-decoding accuracies in Fig. 3b. The CP decoding accuracy (as in Fig. 3b) is shown for comparison in red. (**b**) Same display as Fig. 3b (5-CSWM DMTS data only, red) but showing equivalently determined WM decoding accuracies without including CP variables as predictors (black). Decoding accuracies were still significantly higher than what was achieved by classifiers trained on control datasets with shuffled labels (grey, $P < 10^{-30}$ and $P < 0.002$ for classifiers trained on connectivity or local data, respectively; *t*-test for each connection, not indicated), but are also lower than the accuracies obtained if CP variables are included (red) for all connections and regions (uncorrected *t*-test, indicated by black asterisks). Except for accuracies for MD-derived predictors, the rank order of decoding accuracies across connections/regions remained similar, however. (**c**) Same analysis as in (b) but for the T-maze (5 s delay protocol) and hence in relation to main Fig. 3c. Note that, for several connections/regions, prediction accuracies remained roughly equal if CP predictor variables are omitted (black; uncorrected *t*-test, indicated by black asterisks). Also, decoding accuracies achieved without CP variables were significantly higher than those achieved with classifiers trained the same data but with shuffled labels in all cases (dotted line, $P < 10^{-40}$, *t*-tests, not indicated). Numbers in coloured ovals indicate the rank of the prediction accuracies achieved on average by using data from the respective connection or region. Ranks have been generated from pairwise comparisons with Tukey post-doc tests conducted after significant effects of connection/region in one-way ANOVAs ($P < 0.0001$ in all cases); connections/regions that were not significantly different from each other were assigned the same rank. ns, $P > 0.1$, * $P < 0.05$, ** $P < 0.01$, *** $P < 0.001$. Shaded regions represent s.e.m.

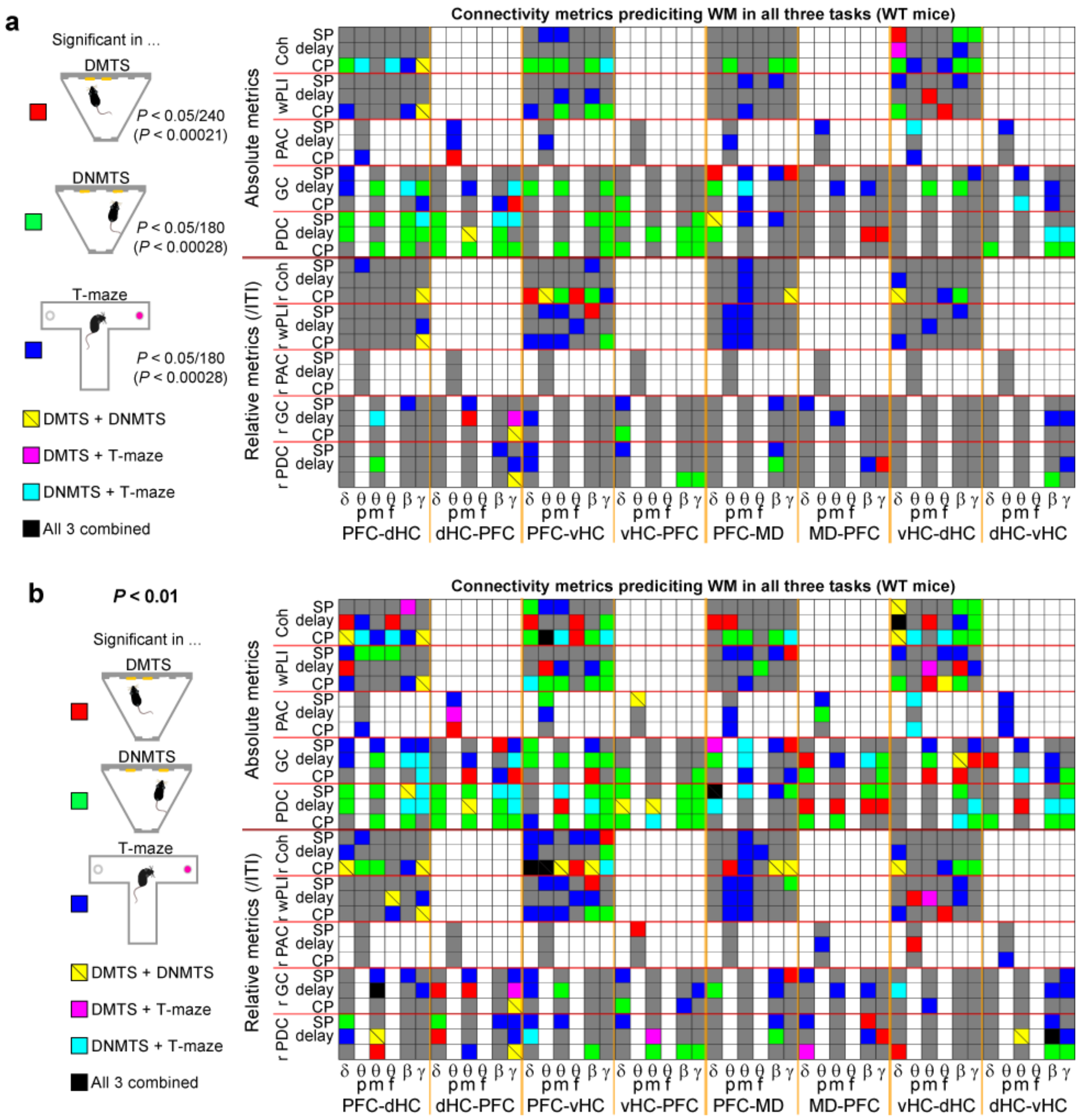


**Supplementary Figure 7 | Individual connectivity measures predicting WM choice across rodent tasks.** (**a-b**) Same display as in main Fig. 4a but for higher (less conservative) *P*-value thresholds: (**a**) Bonferroni-adjustment using the number of parameters per connection (240 for 5-CSWM DMTS, 180 for the other two tasks). (**b**) Use of 0.01 as a fixed threshold. Matrix showing all connectivity predictor variables that contributed to the connection-based classifiers shown in Fig. 3b-c for the three rodent WM tasks (WT only) and were significantly associated with WM-performance according to their weight and differences between correct and incorrect CP (see Results). Note that variables from the pre- and post-delay in the 5-CSWM have been combined in single lines.

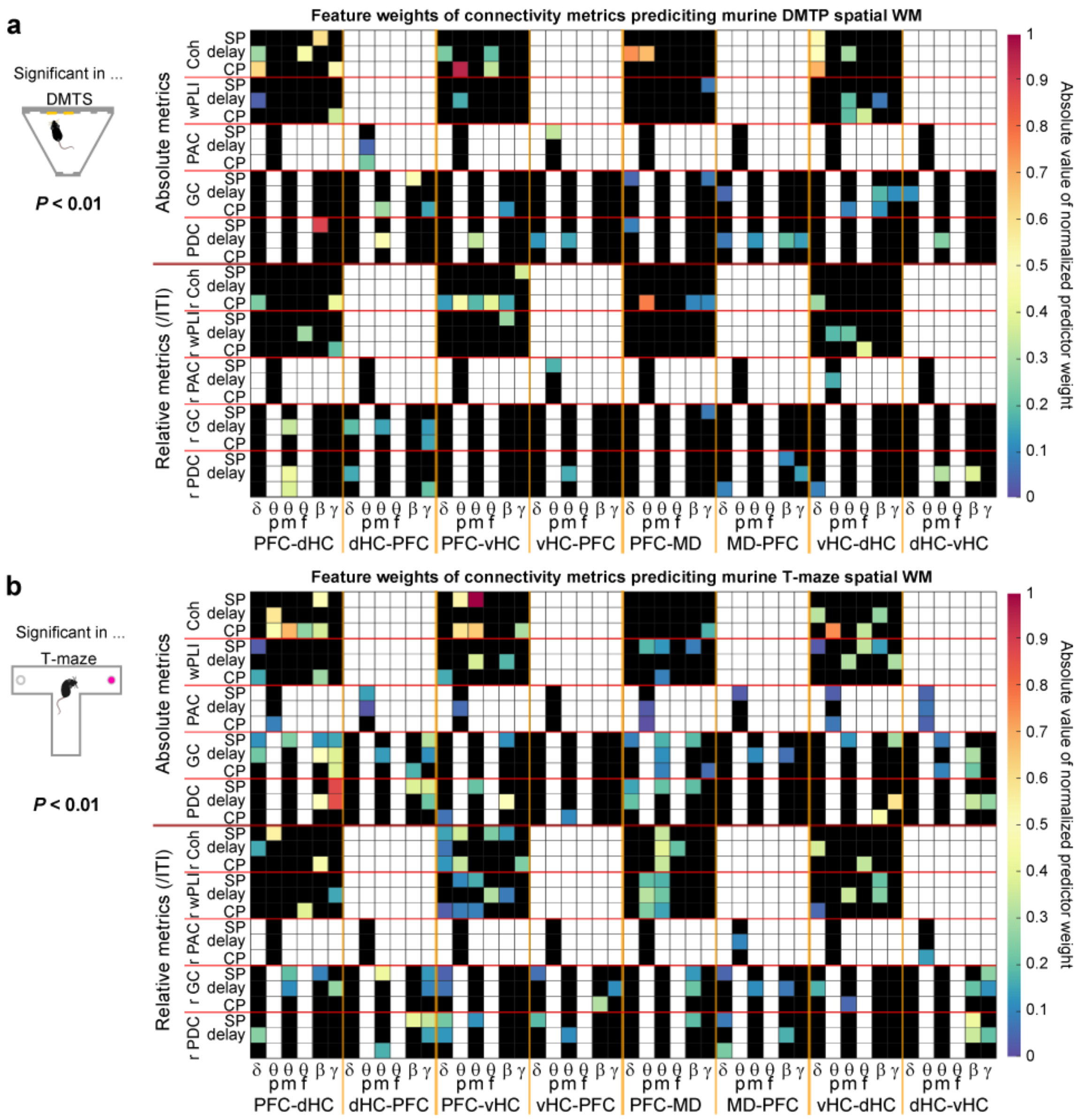


**Supplementary Figure 8 | Predictor weights in mouse 5-CSWM and T-maze.** (**a**,**b**) Matrix of absolute values of the average normalized predictor weights obtained from classifiers calculated for individual connections separately to predict choices in mouse DMTS (a) and T-maze WM (b). Squares corresponding to non-existing metrics in white, squares corresponding to non-significant metrics at a threshold of $P < 0.01$ in black.

a

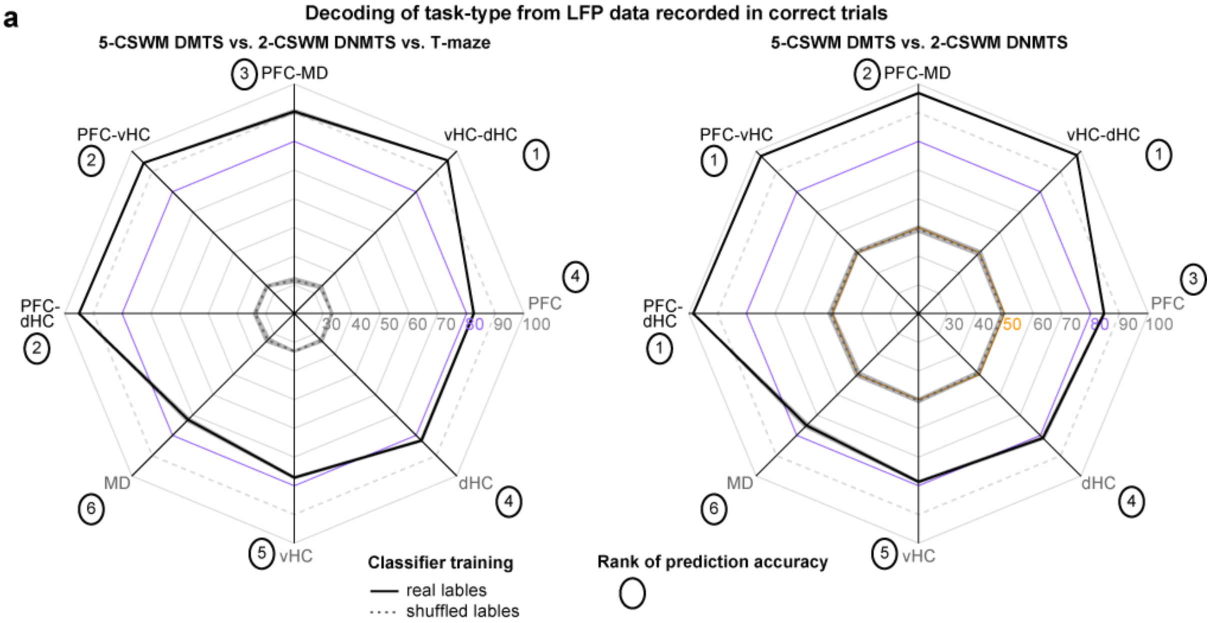


**Supplementary Figure 9 | Prediction of task-type from connectivity and activity data recorded in correct trials.** (**a**) Cross-subject decoding accuracies for predicting the task during which the data was recorded using the connectivity or local activity predictor variables as in the other classifiers (main Fig. 3b-c) albeit only from trials with correct choices. Either all three tasks had to be discriminated from each other (left) or only the two operant tasks (right), resulting in chance levels of 33.3% and 50% respectively. Lines for accuracy levels of 80% and 90% are emphasized by purple colour or dashed appearance, respectively, to aid comparison. Decoding accuracies were significantly higher than what was achieved by classifiers trained on control datasets with shuffled labels (dashed grey lines at chance level, *P* <0.0001; *t*-test for each connection, not indicated). Numbers in coloured ovals indicate the rank of the prediction accuracies achieved on average by using data from the respective connection or region. Ranks have been generated from pairwise comparisons with Tukey post-doc tests conducted after significant effects of connection/region in one-way ANOVAs (*P* < 0.0001 in all cases); connections/regions that were not significantly different from each other were assigned the same rank. Shaded regions represent s.e.m.

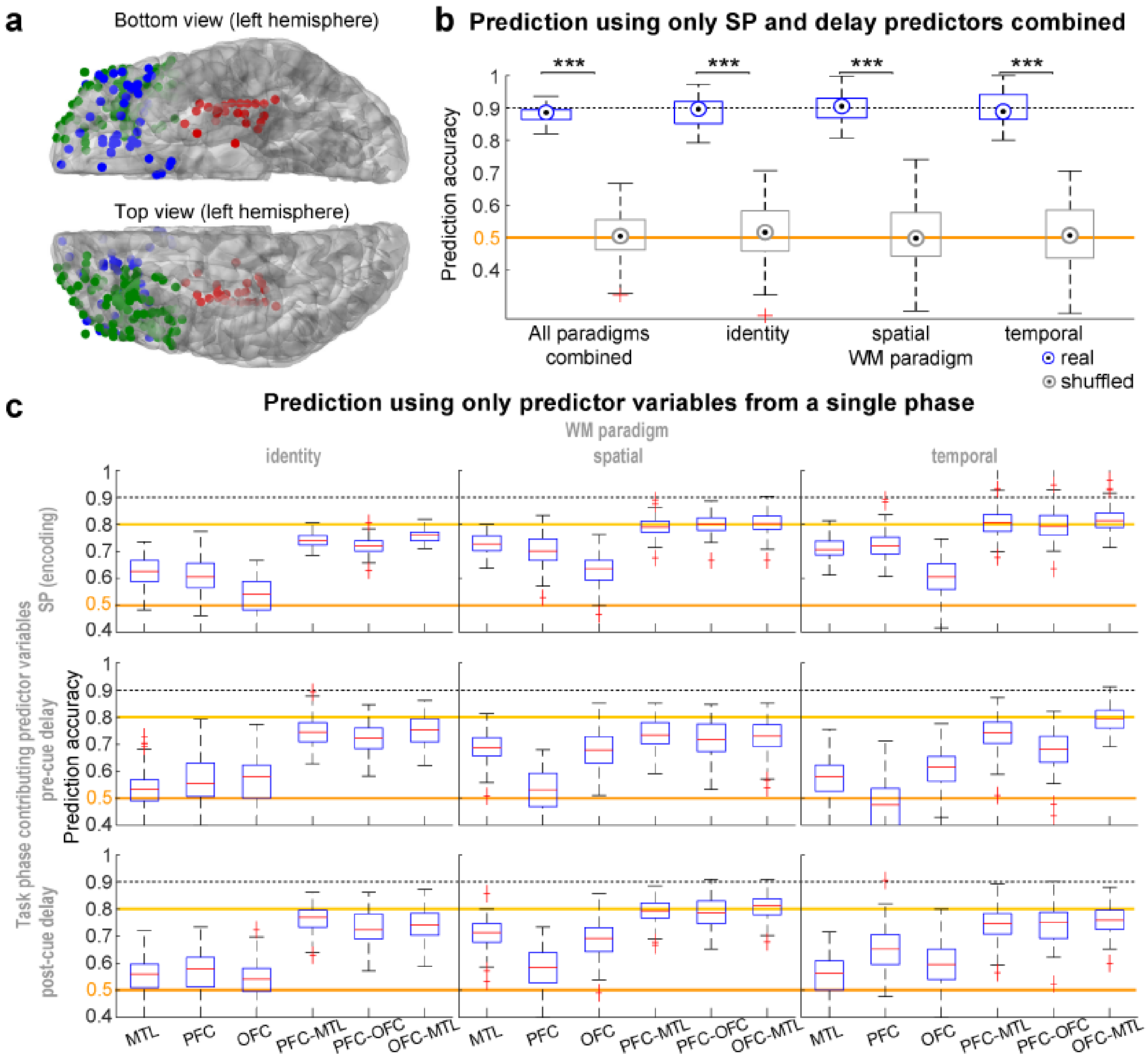


**Supplementary Figure 10 | Decoding accuracy in human WM when using only predictors from the SP and delay. (a)** Positions of electrodes in the 8 subjects included in this analysis viewed from the bottom and from the top of the brain, as indicated; electrode locations are displayed in the left hemisphere irrespective of their actual location in the left or right hemisphere. **(b)** Decoding of human WM performance by including all trials irrespective of sub-task (left) and separately for the identity, spatial and temporal sub-task (as indicated on x-axis), and using variables from all 3 connections and 3 regions combined. In contrast to the primary analysis shown in Fig. 7c-d, here, parameters from the CP and *relative* measures (that proved to carry little predictive weight in the main analysis, see Fig. 8a) were not included as predictor variables. Blue and grey depict performance of classifiers trained on trials with correct or incorrect (shuffled) labels (stars indicate *t*-test comparisons between them). Error bars, data range without identified outliers which are highlighted in red; boxes, range between 25$^{th}$-75$^{th}$ percentile; dot, median. **(c)** Separate classifiers were used to predict WM choice separately for the 3 sub-tasks (named at the top of each subpanel) and for each brain region or connection (named on the x-axis) separately, using exclusively absolute metrics obtained during the during the SP (top, same as main Fig 7f), the pre-cue delay (middle) or the post-cue delay phase (bottom, named on the left). Red lines indicate mean, boxes the 25$^{th}$ and 75$^{th}$ percentile, whiskers indicate data range without outliers, and red crosses indicate outliers. Accuracies of 50% (chance level, orange line), 80% (yellow line) and 90% (dotted line) are indicated to aid comparison in (a,b).

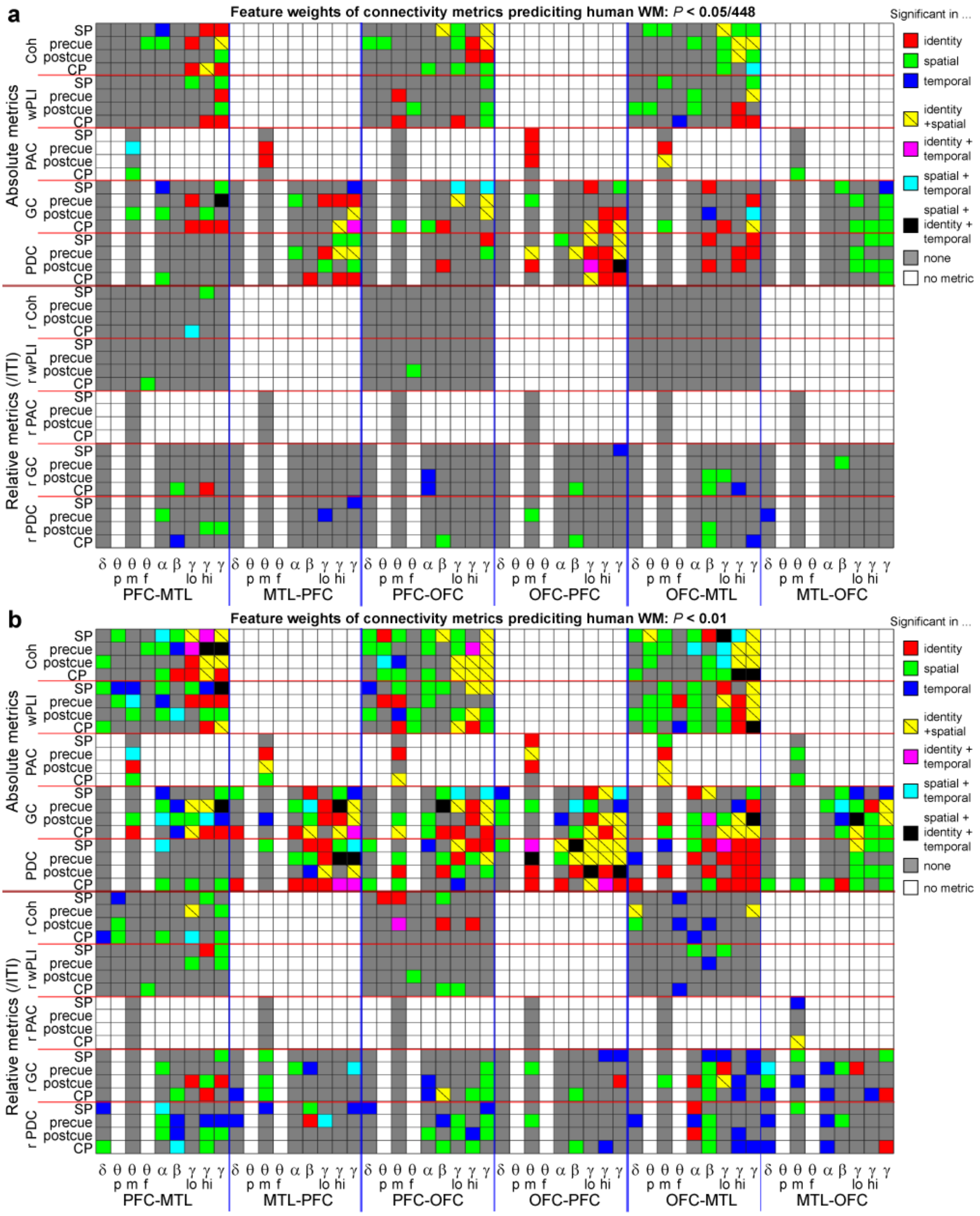


**Supplementary Figure 11 | Individual connectivity measures predicting WM choice in humans selected at lower *P*-value threshold.** (**a-b**) Matrix showing all connectivity predictor variables that contributed to the classifiers for which performance is shown in main Fig. 8a (one classifier using all regional and local variables of all phases, $N$ = 1584, as predictors). Indicated by colour are variables that were significantly associated with WM-performance according to their weight and differences between correct and incorrect CP (see Results) in the paradigms coded by colour (see legend on the right). Same display as in main Fig. 8a, but using less conservative *P*-values: (**a**) Bonferroni-adjustment using the number of predictor variables of each single connection (448), $P < 0.05/448$. (**b**) Standard *P*-value of 0.01, $P < 0.01$. For theta, mean amplitude (m), peak amplitude (p), and frequency of peak (f) are shown, while for all other variables only the mean amplitude is used. The gamma-band contributed three predictors each as this frequency was split into a high- and low-gamma-range in addition to using the whole range (30-100 Hz).

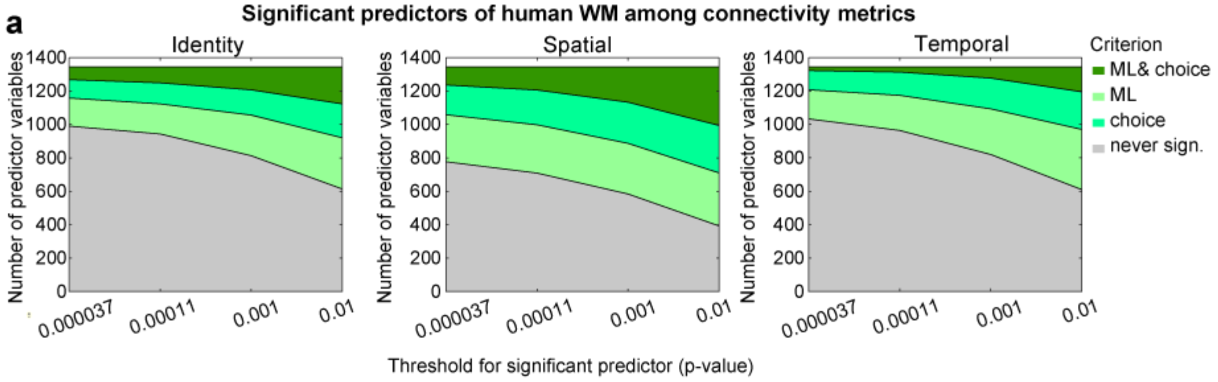


**Supplementary Figure 12 | Number of connectivity measures predicting WM choice in humans in dependence on *P*-value.** (**a**) The number of connectivity variables that are associated with WM-performance as the *P*-value adjustment for the selection of significant predictors is relaxed. The first value in each sub-panel corresponds to the adjustment by the number of *all* connectivity variables ($P < 0.05/1344$), the second value corresponds to the adjustment by number variables in a single connection ($P < 0.05/448$).

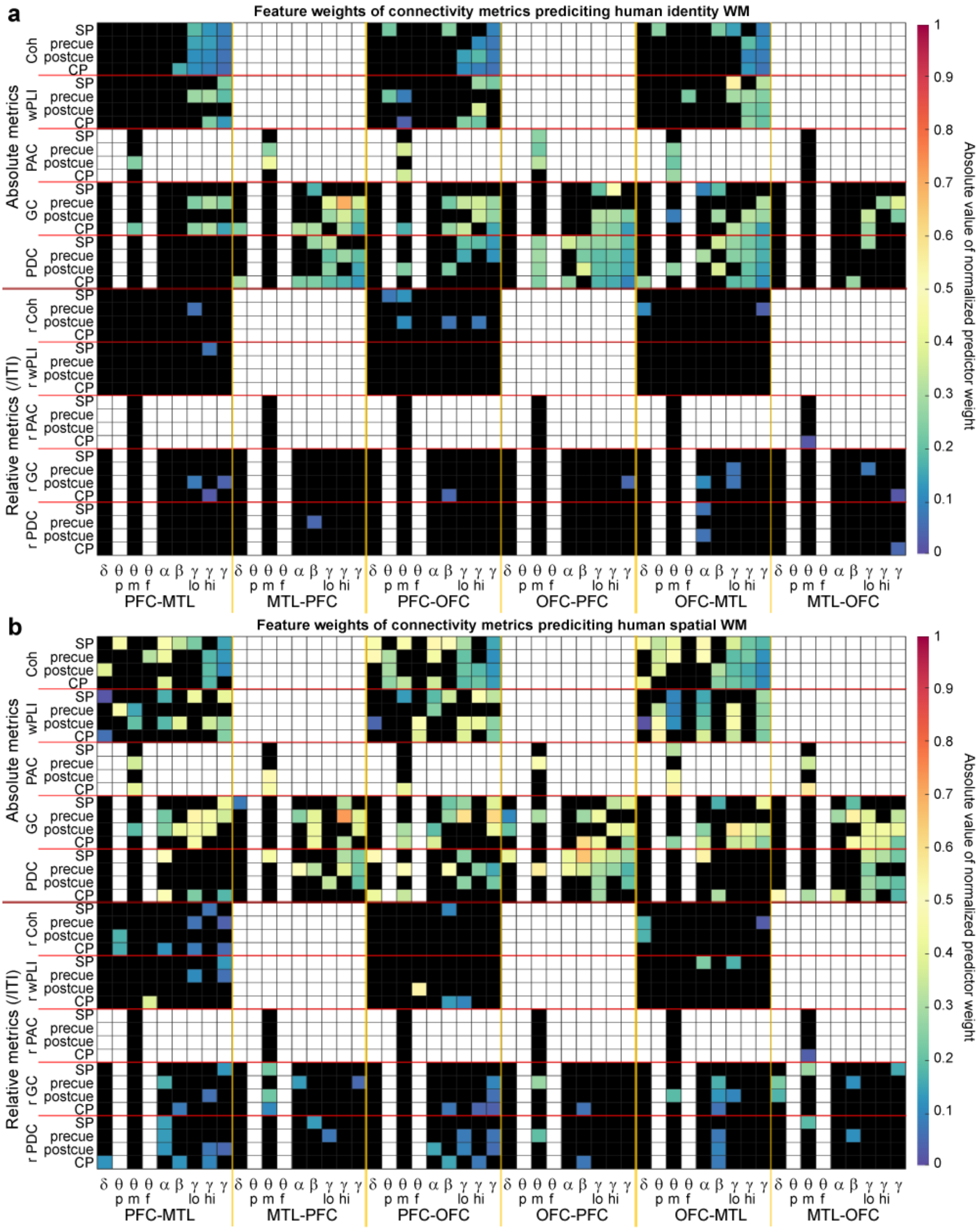


**Supplementary Figure 13 | Predictor weights in human identity and spatial WM tasks.** (**a**,**b**) Matrix of absolute values of average normalized predictor weights obtained from classifiers calculated for all connections and regions combined to predict choices in identityrelated (a) or spatial WM (b). Squares corresponding to non-existing metrics in white, squares corresponding to non-significant metrics at a threshold of $P < 0.01$ in black.

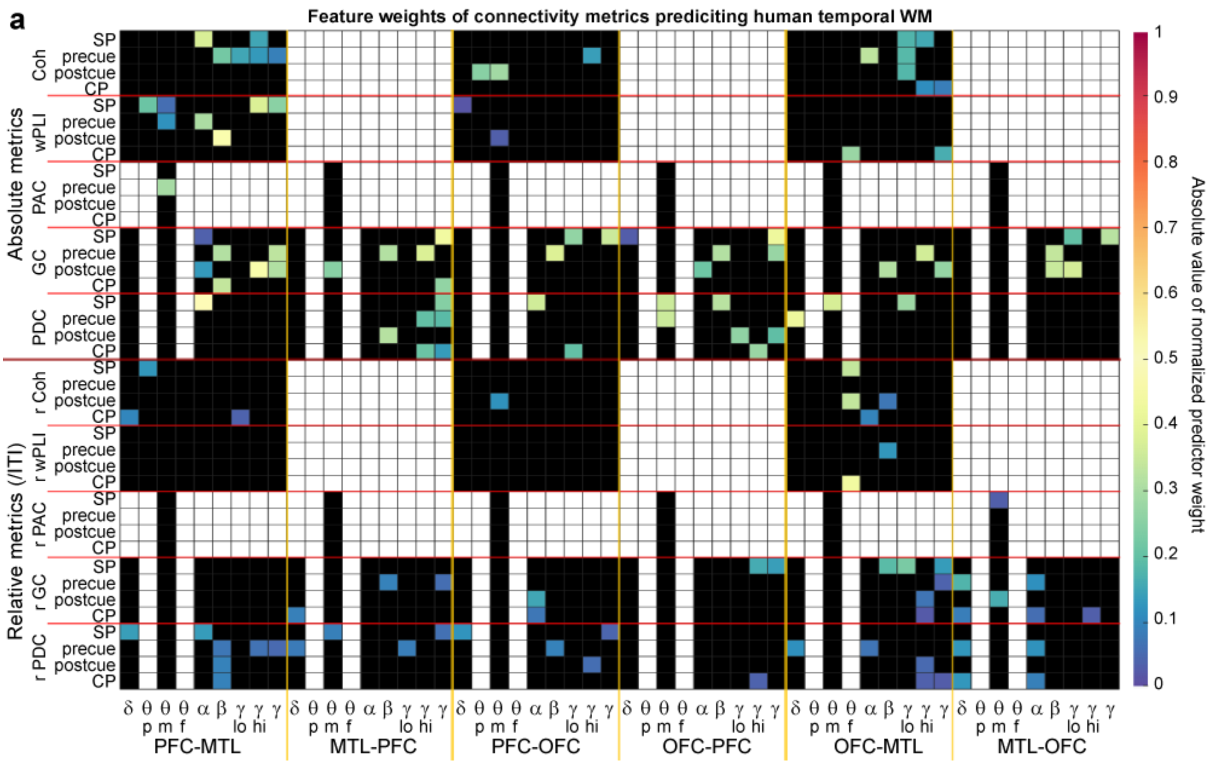


**Supplementary Figure 14 | Predictor weights in human temporal WM task.** (**a**) Same analysis and display as in Supplementary Fig. 13 but depicting predictor weights for classifiers trained on temporal WM.

### Supplementary Tables

| Task | Metric | Frequency | Connection | Ref. |
|---|---|---|---|---|
| T (RA) | Phase synchrony | γ, MUA bursts | dMEC-dCA1 | [1] |
| T (RA) | SPC, coherence | β | PFC(β)-MD (spikes or β) | [2] |
| T (RA) | Coherence | β | PFC-dHC | [3] |
| T (RA) | CFC (PAC) | low γ–θ | PFCγ-vHCθ | [4,5] |
| T (RA) | SPC | γ, SUA | PFC(spikes)-vHC (γ) | [6,7] |
| H (RA) | SPC, Coherence | θ, SUA | PFC(spikes, θ)-vHC (θ) | [8] |
| T (RA) | SPC, Coherence | θ, SUA | PFC(spikes, θ)-vHC (θ) | [4,9,10] |
| T (RA) | SPC, SPC-based directionality (CC) | θ, SUA | Delay: MD→PFC, CP: PFC→MD | [11] |
| T (RA) | CC directionality (lead/lag), coherence | θ, γ | PFC-(Re/Rh)-dHC | [4] |
| 8 (RA) | PDC | θ, α, β | vCA1→PFC, PFC→vCA1 | [12] |
| T (RA) | SPC, Coherence | δ (4 Hz) | VTA (δ)-PFC(δ) | [13] |
| 2-lever operant DNMTS | SPC | θ, SUA | PFC(spikes)-vHC (θ) | [14] |
| Delayed tactile discrimination | SPC, coherence | θ, SUA | PFC(θ)→S1(spikes, θ)- | [15] |

**Supplementary Table 1 | Measures of long-range neural connectivity associated with WM in rodents.** Studies that have attempted to identify an electrophysiological WM correlate in form of coupling of activity in two brain regions are listed, in so far as the association between WM performance and metric have been proven by some analysis, e.g., bivariate correlations, linear regression, or comparison between correct and incorrect trials or between SP and CP phases. Studies that merely noted connectivity and WM differences due to some manipulation independently were not included. **Abbreviations:** T, T-maze; H, H-maze; 8, 8-shaped maze; RA rewarded delayed alternation; SPC, spike-phase-coupling (usually assessed by mean resultant vector length, MRL); CC, cross-correlation of amplitudes to determine temporal leading or lagging of oscillation in one region *vis-a-vis* the other; CFC (PAC) cross-fequency-coupling (phase-amplitude-coupling); PDC, partial directed coherence; MUA, multi-unit activity; SUA, single-unit activity; dMEC, dorsal medial enthorhinal cortex; dCA1/vCA1, dorsal/ventral CA1; dHC/vHC, dorsal/ventral hippocampus;

MD, medio-dorsal thalamus; PFC, medial prefrontal cortex (usually prelimbic area); Re/Rh, Ncl. reuniens / Ncl. rhomboideus of the midline thalamus; VTA, ventral-tegmental area.

| | Accuracy | AUC | F1 | F2 | sensitivity | specificity | cl1-precision | cl1-recall | cl2-precision | cl2-recall |
|---|---|---|---|---|---|---|---|---|---|---|
| **PFC** | 0.66 | 0.66 | 0.66 | 0.65 | 0.65 | 0.67 | 0.68 | 0.65 | 0.64 | 0.67 |
| **dHC** | 0.79 | 0.79 | 0.79 | 0.78 | 0.79 | 0.79 | 0.79 | 0.79 | 0.78 | 0.79 |
| **vHC** | 0.61 | 0.61 | 0.62 | 0.58 | 0.60 | 0.62 | 0.65 | 0.60 | 0.56 | 0.62 |
| **MD** | 0.76 | 0.77 | 0.77 | 0.74 | 0.73 | 0.81 | 0.83 | 0.73 | 0.69 | 0.81 |
| **PFC-dHC** | 0.78 | 0.79 | 0.80 | 0.76 | 0.75 | 0.84 | 0.86 | 0.75 | 0.70 | 0.84 |
| **PFC-vHC** | 0.71 | 0.71 | 0.71 | 0.70 | 0.70 | 0.72 | 0.73 | 0.70 | 0.68 | 0.72 |
| **PFC-MD** | 0.77 | 0.79 | 0.79 | 0.74 | 0.74 | 0.85 | 0.87 | 0.74 | 0.67 | 0.85 |
| **vHC-dHC** | 0.73 | 0.74 | 0.74 | 0.71 | 0.71 | 0.76 | 0.78 | 0.71 | 0.68 | 0.76 |

**Supplementary Table 2 | Classifier parameters for prediction of operant DMTS 5-CSWM in wildtype mice.** Measures of classification performance, the prediction accuracy, the AUC of the receiver operating characteristic (ROC) and the F1-score which is defined as follows: F1 = (2*class1precision * class1recall / (class2precision + class1recall)). Precision is defined as the True positives / True Positive + False Positive for class 1 and as the True negatives / True Negative + False Negative for class 2. Recall is defined as True Positive / True Positive + False Negative which is equivalent to the sensitivity for class1 and specificity for class2.

| | Accuracy | AUC | F1 | F2 | sensitivity | specificity | cl1-precision | cl1-recall | cl2-precision | cl2-recall |
|---|---|---|---|---|---|---|---|---|---|---|
| **PFC** | 0.77 | 0.77 | 0.77 | 0.76 | 0.75 | 0.79 | 0.80 | 0.75 | 0.74 | 0.79 |
| **dHC** | 0.87 | 0.87 | 0.87 | 0.86 | 0.85 | 0.89 | 0.89 | 0.85 | 0.84 | 0.89 |
| **vHC** | 0.71 | 0.72 | 0.73 | 0.69 | 0.69 | 0.74 | 0.77 | 0.69 | 0.65 | 0.74 |
| **MD** | 0.73 | 0.74 | 0.74 | 0.72 | 0.72 | 0.75 | 0.77 | 0.72 | 0.69 | 0.75 |
| **PFC-dHC** | 0.70 | 0.71 | 0.73 | 0.67 | 0.67 | 0.75 | 0.80 | 0.67 | 0.61 | 0.75 |
| **PFC-vHC** | 0.63 | 0.63 | 0.65 | 0.61 | 0.62 | 0.65 | 0.68 | 0.62 | 0.58 | 0.65 |
| **PFC-MD** | 0.67 | 0.68 | 0.69 | 0.63 | 0.65 | 0.71 | 0.76 | 0.65 | 0.58 | 0.71 |
| **vHC-dHC** | 0.65 | 0.66 | 0.67 | 0.63 | 0.64 | 0.68 | 0.71 | 0.64 | 0.59 | 0.68 |

**Supplementary Table 3 | Classifier parameters for prediction of operant DNMTS 2-CSWM in wildtype mice.** Displays same as in Supplementary Table 1 but for the operant DNMTS task.

| | Accuracy | AUC | F1 | F2 | sensitivity | specificity | cl1-precision | cl1-recall | cl2-precision | cl2-recall |
|---|---|---|---|---|---|---|---|---|---|---|
| **PFC** | 0.63 | 0.63 | 0.62 | 0.63 | 0.64 | 0.62 | 0.61 | 0.64 | 0.64 | 0.62 |
| **dHC** | 0.70 | 0.73 | 0.75 | 0.64 | 0.65 | 0.82 | 0.88 | 0.65 | 0.53 | 0.82 |
| **vHC** | 0.67 | 0.67 | 0.66 | 0.67 | 0.68 | 0.67 | 0.66 | 0.68 | 0.68 | 0.67 |
| **MD** | 0.77 | 0.82 | 0.81 | 0.71 | 0.70 | 0.94 | 0.96 | 0.70 | 0.58 | 0.94 |
| **PFC-dHC** | 0.87 | 0.90 | 0.89 | 0.85 | 0.80 | 1.00 | 1.00 | 0.80 | 0.74 | 1.00 |
| **PFC-vHC** | 0.82 | 0.83 | 0.82 | 0.82 | 0.82 | 0.83 | 0.83 | 0.82 | 0.82 | 0.83 |
| **PFC-MD** | 0.84 | 0.88 | 0.86 | 0.80 | 0.77 | 1.00 | 1.00 | 0.77 | 0.68 | 1.00 |
| **vHC-dHC** | 0.83 | 0.85 | 0.84 | 0.81 | 0.79 | 0.91 | 0.91 | 0.79 | 0.75 | 0.91 |

**Supplementary Table 4 | Classifier parameters for prediction of T-maze SWM in wildtype mice.** Displays same as in Supplementary Table 1 but for T-Maze.

| | **Parameters of training and baseline stages** | | | | | |
|---|---|---|---|---|---|---|
| Stage | SP-SD, s | CP-SD, s | Pre-delay, s | Post-delay, s | Reward (μl) | CP configurations |
| 1 | 20 | 20 | 0 | 2 | 20 | 1-3, 2-4, 3-5 |
| 2 | 20 | 20 | 0 | 2 | 10 | 1-3, 2-4, 3-5 |
| 3 | 20 | 20 | 0 | 2 | 0 | 1-2, 2-3, 3-4, 4-5 |
| 4 | 20 | 20 | 0 | 2 | 10 | 1-2, 1-3, 1-4, 2-3, 2-4, 2-5, 3-4, 3-5, 4-5 |
| 5 | 10 | 5 | 0 | 2 | 10 | 1-2, 1-3, 1-4, 2-3, 2-4, 2-5, 3-4, 3-5, 4-5 |
| 6 | 2 | 5 | 0 | 2 | 10 | 1-2, 1-3, 1-4, 2-3, 2-4, 2-5, 3-4, 3-5, 4-5 |
| | **Parameters of challenge stages** | | | | | |
| Delay 5s | 10 | 5 | 5 | 2 | 10 | As baseline |
| Delay 10s | 10 | 5 | 10 | 2 | 10 | As baseline |
| Distraction | 10 | 5 | 0 | 2 (Distract.) | 10 | As baseline |
| Attention | 1 | 5 | 0 | 2 | 10 | As baseline |
| Combined | 1 | 5 | 5 | 2 | 10 | As baseline |

**Supplementary Table 5. 5-CSWM training stages.** Note that, on *all* stages, the *post-delay* is 2 s, the *CP reward* is 60 μl, the *limited hold time* - that is the time in which a response is registered - exceeds the SD by 1 s, the *ITI* is 5 s, and the time-out time after incorrect responses or omissions is *5 s*. The CP configurations indicate the two holes of the 5-choice wall that can be illuminated; note that the actual configurations when taking into account the correct hole is double than what is stated (i.e., 2-4 is different from 4-2, which is not listed). Stage 4 served a pre-surgery baseline and post-surgery training stage without tether. Stage 5 was only conducted with simultaneous recordings (or tether) and served as baseline for all challenge conditions conducted with simultaneous recordings, except that – for habituation to shorter stimulus intervals – the last challenges (with SP-SD 1 s) had their own baseline as reference, featuring an SP-SD of 2 s: stage 6.

| | # of electrodes | # of mice | Number of trials in ... | | | |
|---|---|---|---|---|---|---|
| | | | 5-CSWM, 1s SD + 5s delay | DNMTS, baselline | T-maze, 5s | T-maze, 30s |
| PFC electrodes | 19 | 12 | 758 | 1431 | 732 | 674 |
| dHC electrodes | 7 | 7 | 464 | 1320 | 448 | 408 |
| vHC electrodes | 12 | 9 | 584 | 1700 | 544 | 480 |
| MD electrodes | 4 | 7 | 236 | 694 | 246 | 224 |
| PFC-dHC | 7 | 7 | 464 | 1320 | 448 | 408 |
| PFC-vHC | 9 | 9 | 584 | 1700 | 544 | 480 |
| vHC-dHC | 6 | 6 | 400 | 1172 | 318 | 334 |
| MD-PFC | 4 | 4 | 236 | 694 | 246 | 224 |

**Supplementary Table 6. Number of electrodes, connections, mice and trials in mouse experiments.**
